# Genetic pathways regulating the longitudinal acquisition of cocaine self-administration in inbred and recombinant inbred mice

**DOI:** 10.1101/2022.11.10.516062

**Authors:** Arshad H. Khan, Jared R. Bagley, Nathan LaPierre, Carlos Gonzalez-Figueroa, Tadeo C. Spencer, Mudra Choudhury, Xinshu Xiao, Eleazar Eskin, James D. Jentsch, Desmond J. Smith

## Abstract

To identify genetic pathways for addiction, we analyzed intravenous self-administration of cocaine or saline in a panel of 84 inbred and recombinant inbred mouse strains over 10 days. We integrated the behavior data with RNA-Seq data from the medial frontal cortex and nucleus accumbens from 41 strains. The self-administration of cocaine and saline showed distinct genetic bases. We maximized power to map loci for cocaine intake by using a linear mixed model to account for this longitudinal phenotype while correcting for population structure. A total of 15 unique significant loci were identified in the genome-wide association study (GWAS). A transcriptome-wide association study (TWAS) highlighted the *Trpv2* ion channel as a key locus for cocaine self-administration from the GWAS. In addition, 17 genes supplementary to the GWAS were identified including *Arhgef26, Slc18b1* and *Slco5a1*. We found numerous instances where alternate splice site selection or RNA editing altered transcript abundance. Our work emphasizes the importance of *Trpv2*, a known cannabinoid receptor, for the response to cocaine as well as identifying further relevant loci.

## Introduction

Cocaine use disorders are a significant economic and health burden. In the US, a total of 2 million people use the drug once a month or more and greater than 850,000 individuals reach the criteria for cocaine dependence (Deak and Johnson, 2021; Fernàndez-Castillo et al., 2022; Palmer et al., 2021; Pierce et al., 2018). The rate of deaths due to overdose in 2018 were estimated at 4.5 per 100,000 standard population (Hedegaard et al., 2020).

Cocaine acts by blocking the reuptake transporters for dopamine, serotonin and norepinephrine in presynaptic nerve terminals, thus increasing the concentrations of these neurotransmitters in the synaptic cleft. The rewarding effects of cocaine is largely mediated by increased dopaminergic neurotransmission in the limbic system, in particular the nucleus accumbens (NAc) as well as the prefrontal cortex (Fernàndez-Castillo et al., 2022; Pierce et al., 2018).

Cocaine addiction is caused by the interplay between multiple environmental and genetic factors, and is hence a complex trait. The broad sense heritability (H^2^) for cocaine use is ~0.32–0.79 and the additive heritability (h^2^) for cocaine dependence is ~0.27–0.30. There is evidence of overlap between the genetic risk factors for cocaine addiction and other addictive drugs, in particular cannabis (Kendler et al., 2007).

Properly powered genome-wide association studies (GWASs) of cocaine addiction in humans await ascertainment of adequate population sizes, likely tens to hundreds of thousands of individuals. Hurdles in obtaining sufficient study numbers include difficulties in recruiting cocaine-dependent individuals, gene-environment interactions and population and phenotypic heterogeneity. One genome-wide significant variant for cocaine use disorder has been identified in the *FAM53B* gene (Gelernter et al., 2014). Analysis of the same data using a gene-based test identified *C1QL2*, *STK38*, *KCTD20* and *NDUFB9* as addiction genes, while another meta-analysis identified *HIST1H2BD* (Cabana-Domínguez et al., 2019; Huggett and Stallings, 2020a, 2020b). One recent GWAS employed gene-environment interactions to identify 13 significant genes, while another recent study found two significant loci associated with time from cocaine use to dependence (Sherva et al., 2021; Sun et al., 2020).

Genetic studies in mice can provide useful insights into cocaine addiction, since careful control of environment and behavioral endpoints can be easily accomplished compared to humans. One investigation evaluated cocaine self-administration in 39 strains of recombinant inbred BXD mice. A cocaine self-administration quantitative trait locus (QTL) was found to harbor a *trans* expression QTL (eQTL) for *Fam53b* (Dickson et al., 2016). Another study using BXD mice showed that poor reversal learning, which indicates a lack of inhibitory control, was associated with greater cocaine self-administration (Cervantes et al., 2013).

Further evidence for a genetic basis of cocaine use was the observation that an acute dose of the drug caused significant differences in locomotor activity across 45 inbred mouse strains (Wiltshire et al., 2015). Divergence in sensitization to cocaine was also found using 51 genetically diverse collaborative cross strains and their inbred founders (Schoenrock et al., 2022). An elegant study using two closely related substrains of the C57BL/6 mouse strain revealed that a missense mutation in *Cyfip2* results in altered sensitization to cocaine (Kumar et al., 2013).

The hybrid mouse diversity panel (HMDP) is a collection of ~30 inbred and ~70 recombinant inbred mouse strains that can be used for association mapping of complex traits including behavior (Ghazalpour et al., 2012, 2012; Lusis et al., 2016; Park et al., 2011a). The inbred strains have a large number of breakpoints that facilitates high resolution genetic mapping, while the recombinant inbred strains increase statistical power. Because the HMDP is genetically stable it is possible to layer multiple phenotypes on the panel, providing ever more powerful insights.

We have previously used the HMDP to evaluate cocaine and saline intravenous self-administration (IVSA) over a ten day testing period (Bagley et al., 2022). The panel showed high phenotypic diversity in cocaine and saline IVSA, consistent with a genetic basis for these traits. Genetic mapping revealed significant loci for cocaine IVSA on Chromosomes 3 and 14. Massively parallel RNA sequencing (RNA-Seq) of the nucleus accumbens (NAc) and medial frontal cortex (mFC) provided *cis* expression QTLs (eQTLs) that could be employed to narrow down candidate genes for cocaine self-administration. However, the power of the study was not fully realized since the genome-scans used binned data from five sequential sets of two individual days, rather than exploiting the longitudinal nature of the datasets.

In this study, we used a linear mixed model to analyze the same dataset and identify loci that affect longitudinal cocaine IVSA while correcting for population structure. We evaluated the RNA-Seq data for *cis* and *trans* QTLs affecting transcript, spliceform and editing abundance. Splicing and editing events that influenced transcript abundance were identified. We then used transcriptome-wide association studies (TWASs) to combine the results of the behavioral GWASs with the RNA-Seq data, providing confirmatory support for genes underlying cocaine use while also suggesting additional genes.

## Materials and Methods

### Cocaine intravenous-self administration

A total of 479 and 477 mice from 84 strains of the HMDP were used for cocaine and saline IVSA respectively, as described earlier (Bagley et al., 2022). A total of 32 inbred and 52 recombinant inbred strains were evaluated. Animals were acquired from the Jackson Laboratory (Bar Harbor ME) with an indwelling jugular catheter. Target numbers were 3 males and 3 females for each strain for each of the two infusates. The actual number per strain was 5.7 ± 0.1 s.e.m. for both cocaine and saline, exceeding the 5 animals calculated to provide 80% power to identify a QTL with an effect size of 10% in the 100 strains of the HMDP (Bennett et al., 2010). There were close to equal numbers of males and females within each strain and infusate (50.4 ± 0.7 % males). The age of the mice was 11.3 ± 0.07 weeks.

Mice were evaluated for cocaine or saline IVSA in 10 consecutive daily sessions as described (Bagley et al., 2022). Animals were confronted with two levers in the testing chambers, one of which gave an infusion of cocaine or saline, the other of which did not. A time-out period of 20 s occurred after an infusion, during which active lever presses were recorded but no infusion was given. Testing continued until 65 infusions were administered or 2 h elapsed, whichever came first. Four endpoints were analyzed; number of infusions, number of active lever presses, percentage of active lever presses and number of inactive lever presses.

### Genome-wide association studies of IVSA

Genetic mapping of loci for cocaine and saline IVSA was performed for individual days using a linear mixed model implemented by `FaST-LMM` to correct for population structure via a kinship matrix (Bagley et al., 2022; Lippert et al., 2011). Covariates included sex, active lever (left or right), testing chamber, cohort and age. Behavioral data was normalized using the rank-based inverse normal transformation (Blom transformation) (Mangiafico, 2022). Genome-wide significance thresholds were obtained from permutation, *P* < 4.1 × 10^−6^, as described (Bennett et al., 2010; Hasin-Brumshtein et al., 2016). Single nucleotide polymorphism (SNP) genotypes were obtained from the mouse diversity array (Rau et al., 2015). After removing SNPs with minor allele frequency < 5% or missing genotype frequency > 10%, 340,097 SNPs remained for mapping. Coordinates are from mouse genome build GRCm38/mm10 (Kent et al., 2002).

To increase power to detect significant loci for cocaine IVSA, we used `GMMAT` to implement a linear mixed model that treated the phenotypes as longitudinal traits over the 10 days of testing (Chen et al., 2016). This model included fixed and random effects of day of testing and also corrected for population structure using a kinship matrix. Covariates were as described above for `FaST-LMM`. The Wald test was used to determine significance because of its increased power compared to the score test in linear models (Demidenko, 2020). One genome scan took ~4 weeks on a computer cluster.

### Heritability

Behavioral covariates were evaluated using a linear mixed model implemented in `lme4` with day of assay, sex, chamber number, active lever and age as fixed effects and strain as a random effect (Bates et al., 2015). Broad sense heritability (H^2^) was calculated using the same model, but with day of assay omitted to evaluate H^2^ on individual days. Additive heritability (h^2^) was calculated using the `heritability` package (Kruijer and White, 2019).

### RNA-Seq

NAc (core and shell) and mFC were harvested from all mice 24 h after their final test session (Bagley et al., 2022). RNA-Seq was performed on 41 strains exposed to either cocaine or saline, consisting of 28 inbred and 13 recombinant inbred strains. In four cocaine and four saline strains, RNA-Seq was done using individual samples, yielding a total of 6 samples composed of 3 males and 3 females for each infusate and brain region. For the remaining strains, samples of the same sex were pooled, yielding a total of 2 samples for each infusate and brain region. A total of 392 samples were analyzed using RNA-Seq with 75 bp paired-ends. Sequencing depth per strain for each brain region was 72.6 ± 1.1 million (M) reads for cocaine and 73.0 ± 1.1 M for saline. Reads were mapped to the mouse genome using STAR aligner (Bagley et al., 2022; Dobin and Gingeras, 2016; Hasin-Brumshtein et al., 2016).

### GWAS of transcript abundance

Transcripts that had ≥ 6 reads and transcripts per million (TPM) > 0.1 in at least 20% of samples for each infusate (cocaine or saline) and brain region (NAc or mFC) were selected for analysis (Hasin-Brumshtein et al., 2016; The GTEx Consortium, 2020). A total of 21,118 ± 41 transcripts remained for analysis, averaged over the two brain regions and infusates. Conditional quantile normalization was used to normalize the data (Hansen et al., 2012). Expression quantitative trait loci (eQTLs) were mapped using `FaST-LMM` with covariates of sex and sequencing batch. *Cis* eQTLs were defined as residing within 2 Mb of the regulated gene, with genome-wide significance thresholds of *P* < 1.4 × 10^−3^ for *cis* eQTLs and 6 × 10^−6^ for *trans* (Hasin-Brumshtein et al., 2016). Pairs of QTLs, whether behavioral or molecular, were defined as coincident if they were located within 2 Mb of each other.

### GWAS of splicing

Exons were selected with ≥ 5 reads in all samples for each infusate and brain region. The exon in each gene with the highest standard deviation of percentage spliced in (psi, or ψ) between individuals was then selected for further study (Kahles et al., 2018). Only exons with non-zero standard deviation that were also included in the transcript analysis were retained. A total of 9,436 ± 59 exons remained for analysis, averaged over the two brain regions and infusates. Values of ψ were quantile normalized and used to map QTLs for alternate splicing (ψQTLs) as described for transcripts, above.

### GWAS of RNA editing

We quantified RNA editing sites by aligning RNA-Seq reads using hisat2 v.2.0.4 with default parameters (Kim et al., 2019). Unmapped reads were realigned using a pipeline to resolve mapping of hyper-edited reads (Porath et al., 2014; Tran et al., 2019). RNA editing sites were then obtained from the REDIportal database and downstream processing steps performed as described (Bahn et al., 2012; Lee et al., 2013; Picardi et al., 2017; Zhang and Xiao, 2015).

Editing sites were retained for further analysis if ≥ 10% of samples for each infusate and brain region had data. A total of 5,266 ± 181 editing sites were analyzed, averaged over the two brain regions and infusates. We quantile normalized editing ratios (ϕ) and used `FaST-LMM` to map QTLs regulating RNA editing (ϕQTLs). All editing sites were A to I.

### Regulation of transcripts, spliceforms and editing sites

To ensure that we identified expression changes independent of genetic background, we used a linear mixed model implemented in `lme4qtl` (Ziyatdinov et al., 2018). Fixed effects were brain region, sex, infusate, sex/infusate interaction and batch. For splicing and editing, transcript expression level was added as an additional fixed effect. Random effects used a kinship matrix to account for population structure and to correct for regulatory effects due solely to genetic background.

### Transcriptome-wide association studies

Transcriptome-wide association studies (TWASs) were performed using FUSION and FOCUS software (Gusev et al., 2016; Mancuso et al., 2019). Significance thresholds for FUSION were established using *P* < 0.05, Bonferroni corrected for the number of genes tested. FOCUS genes were considered significant if the posterior inclusion probability (pip) > 0.8.

### eCAVIAR

To find SNPs that co-regulated *cis* eQTLs and behavioral loci with the highest colocalization posterior probability (CLPP), we used eCAVIAR to evaluate markers within 200 SNPs of the *cis* eQTL (Hormozdiari et al., 2016). A CLPP > 0.01 is considered the support threshold for a co-regulating SNP.

### Data availability

The sequencing data generated in this study can be downloaded from the NCBI BioProject database (https://www.ncbi.nlm.nih.gov/bioproject/) under accession number PRJNA755328. Data and code is also available from figshare (https://figshare.com/; doi: 10.6084/m9.figshare.21539487)

## Results

### Cocaine and saline self-administration

As described previously, we evaluated 84 strains of the HMDP for cocaine (479 mice) or saline IVSA (477 mice) over a 10 day testing period (Supplementary Table S1) (Bagley et al., 2022). Animals could press either of two levers in the testing chamber; one caused the infusate (cocaine or saline) to be delivered, the other was inactive. Four behavioral endpoints were evaluated; number of infusions, active lever presses, percent active lever presses and inactive lever presses.

### Behavioral covariates

To evaluate influences on the behavioral endpoints independent of genetic background, we used a linear mixed model with fixed effects of chamber identity, active lever (left vs right), age, sex and testing day, together with a random effect of strain (Supplementary Figures S1-S4).

There was a significant effect of testing chamber on all behavioral measures for both cocaine (χ^2^ = 172, df = 45, *P* < 2.2 × 10^−16^, inactive lever presses, least significant end points quoted) and saline (χ^2^ = 177, df = 46, *P* < 2.2 × 10^−16^, percent active lever presses) (Supplementary Figure S1). Testing chambers were assigned non-randomly, to minimize the evaluation of multiple mice from the same strain in the same chamber (Bagley et al., 2022). The significant effect of testing chamber may be partly due to the fact that not all chambers received a representative balance of strains.

Further, due to the large sample size, covariates were apt to be significant even for small effects. For example, there was significantly higher self-administration of both cocaine (t[1, 4687] = 6.4, *P* = 2.3 × 10^−10^, Kenward-Roger df, active lever presses) and saline (t[1, 4651] = 2.6, *P* = 0.011, infusions) when the active lever was on the left compared to the right (Supplementary Figure S2). However, the difference in active lever presses for cocaine was only 6.5 ± 1.3 (35.6 ± 1.0 for left; 29.1 ± 0.9 for right) and 0.8 ± 0.6 for saline infusions (15.7 ± 0.4 for left; 15.0 ± 0.4 for right). Inactive lever presses were significantly higher when the lever was on the right (cocaine, t[1, 4708] = 4.8, *P* = 1.7 × 10^−6^; saline, t[1, 4700] = 7.3, *P* = 4.6 × 10^−13^).

Animal age had varied effects on the behavioral endpoints, both significant and non-significant (Supplementary Figure S3). There was no significant effect of sex on cocaine or saline self-administration, though males showed significantly higher presses of the inactive lever for both cocaine (t[1, 4706] = 5.2, *P* = 1.7 × 10^−7^) and saline (t[1, 4699] = 2.6, *P* = 9.1 × 10^−3^) (Supplementary Figure S4).

There were significant increases in self-administration over the 10 day course of the experiment for both cocaine (t[1, 4657] = 9.0, *P* = 5.1 × 10^−19^, active lever presses) and saline (t[1, 4636] = 6.0, *P* < 1.8 × 10^−9^, percent active lever presses) (Supplementary Figures S5-S6). However, animals working for cocaine showed significantly higher infusions (t[1, 9431] = 19.5, *P* < 2.2 × 10^−16^) and percent active lever presses (t[1, 9464] = 26.2, *P* < 2.2 × 10^−16^) than those responding to saline. There was no significant effect of infusate for active lever presses (t[1, 9446] = 0.08, *P* = 0.93). In contrast, inactive lever presses were significantly greater for saline than cocaine (t[1, 9460] = 4.5, *P* = 7.2 × 10^−6^).

### Differing genetic basis for cocaine and saline IVSA

We found distinct genetic mechanisms for cocaine and saline-taking. Although there were significant correlations between infusates for behavioral measures averaged by strain (cocaine vs saline), there were higher correlations within the same infusate (cocaine vs cocaine and saline vs saline; t = 18.8, df = 2408, *P* < 2.2 × 10^−16^), suggesting distinct behavioral responses to the two regimens (Figure 1A, Supplementary Figure S7).

**Figure 1.**
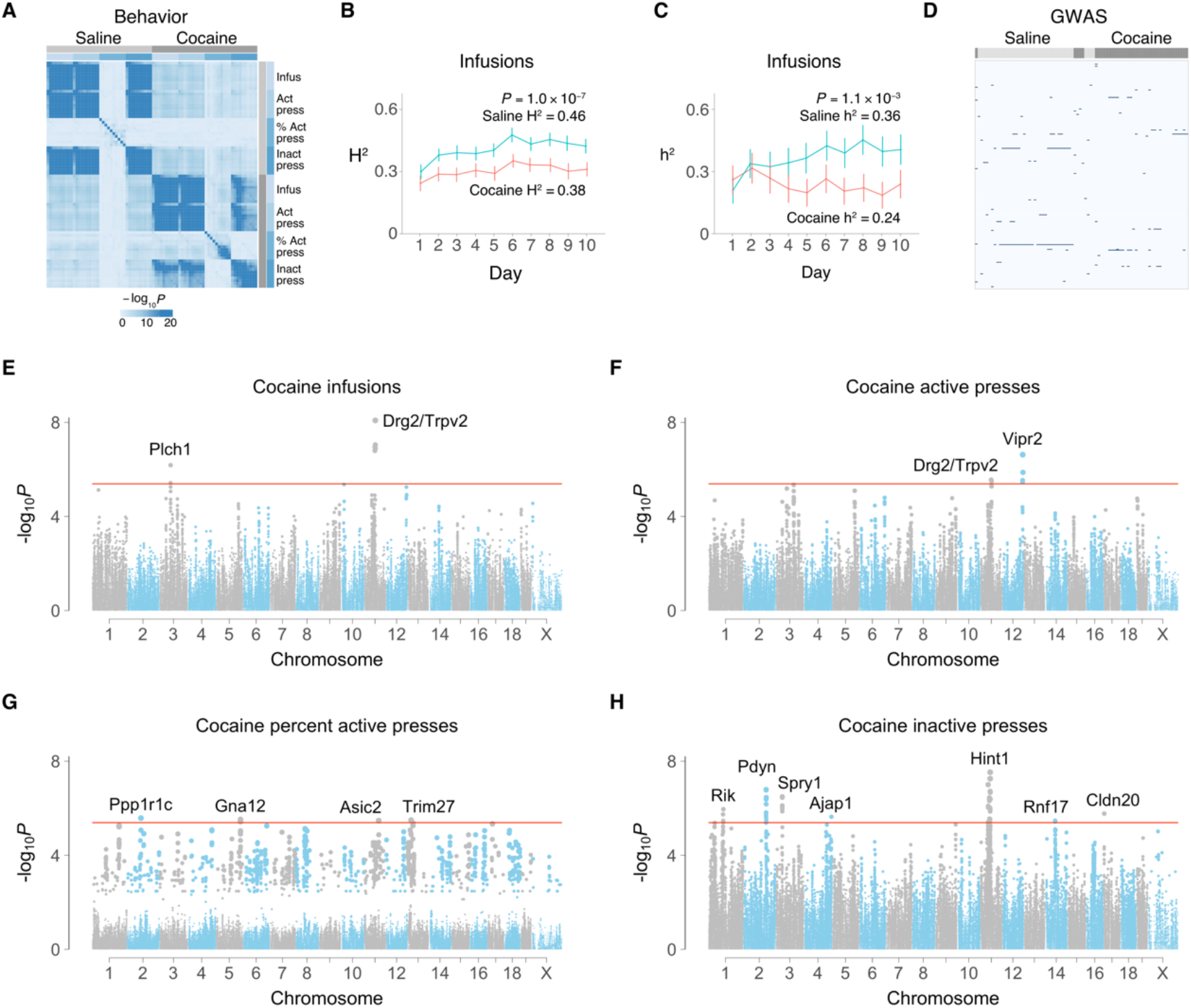
Cocaine and saline IVSA. (**A**) Correlations between behaviors averaged by strain. (**B**) Broad sense heritability (H^2^) for cocaine and saline infusions over 10 days. P value compares cocaine and saline. (**C**) Additive heritability (h^2^). (**D**) Clustering of genome wide association studies (GWASs) for cocaine and saline IVSA. Columns represent each of the ten days for infusions, percent active lever presses, active lever presses and inactive lever presses. Rows represent SNPs in genome order. Color represents GWAS −log_10_P value. (**E**) Longitudinal genome scans for cocaine infusions, (**F**) active presses, (**G**) percent active presses, (**H**) inactive presses. Rik, A630001G21Rik. Red horizontal line, family-wise error rate = 5%.

Over the 10 day period, all measures of self-administration showed significant broad sense heritability (H^2^), both for saline (e.g. infusion H^2^ = 0.46 ± 0.001, χ^2^ = 2355, df = 1, P < 2.2 × 10^−16^) and cocaine (infusion H^2^ = 0.38 ± 0.001, χ^2^ = 1756, df = 1, P < 2.2 × 10^−16^) (Figure 1B, Supplementary Figures S7-S8). However, there was significantly higher broad sense heritability for saline compared to cocaine for infusions, active lever presses and inactive lever presses (*P* < 2.7 × 10^−5^, sampling without replacement).

Additive heritability (h^2^) was also significant for all measures of self-administration for both saline (infusion h^2^ = 0.36 ± 0.02, t[1,9] = 17.0, *P* = 3.8 × 10^−8^, one sample t-test) and cocaine (infusion h^2^ = 0.24 ± 0.01, t[1,9] = 18.8, *P* = 1.6 × 10^−8^) (Figure 1C, Supplementary Figure S9). Similar to broad sense heritability, additive heritability was significantly higher for saline than cocaine for infusions, active lever presses and inactive lever presses (infusions, χ^2^ = 10.7, df = 1, *P* =1.1 × 10^−3^). Broad sense and additive heritabilities were roughly consistent with estimates from human populations.

To further explore the genetic basis for cocaine and saline IVSA, we performed a genome-wide association study (GWAS) for each of the ten days using infusions, active lever presses, percentage active lever presses and inactive lever presses (Bagley et al., 2022). A linear mixed model using `FaST-LMM` software was employed to correct for population structure (Lippert et al., 2011). Although none of the GWASs for the individual days exceeded genome-wide significance, clustering of the results showed segregation of the genome scans for cocaine and saline (Figure 1D). Further, there was a significantly higher correlation of GWAS results within infusate (cocaine vs cocaine and saline vs saline) than between (cocaine vs saline) (t[1, 1898] = 29.0, *P* < 2.2 × 10^−16^) (Supplementary Figure S10). Together, these observations indicate different genetic factors for cocaine and saline IVSA.

### Increasing statistical power using a longitudinal analysis

To improve statistical power for GWA, we used `GMMAT` software to run a linear mixed model that can incorporate longitudinal phenotypes (Chen et al., 2016). The day of assay was used as both a fixed and random effect together with fixed effect covariates of age, sex, active lever position (left or right), conditioning chamber and cohort, together with random effects of SNPs to correct for genetic relatedness. A total of 17 significant loci were identified, of which 15 were unique (Table 1, Figures 1E-H).

**Table 1.** Loci for cocaine self-administration

| Chr | SNP | Pos (bp) | *P* | Assay | Gene | Evidence<sup>1</sup> |  |  |  |
| --- | --- | --- | --- | --- | --- | --- | --- | --- | --- |
|  |  |  |  |  |  | Distance (bp)<sup>2</sup> | FUS<sup>3</sup> | FOC<sup>4</sup> | *Cis* eQTL<sup>5</sup> |
| 1 | rs33037178 | 85,670,029 | 1.1E-06 | Inact | *A630001G21Rik* | 77,834 | N | N, mF | N, mF |
| 2 | rs28034317 | 79,739,802 | 2.6E-06 | % Act Press | *Ppp1r1c* | 23,336 |  |  |  |
| 2 | rs27257529 | 129,620,509 | 1.6E-07 | Inact | *Pdyn* | 72,695 |  |  |  |
| 3 | rs30059671 | 38,178,200 | 3.2E-07 | Inact | *Spry1* | -535,928 |  |  |  |
| 3 | rs50587939 | 63,815,060 | 6.6E-07 | Infus | *Plch1* | -17,207 |  |  | mF |
| 4 | rs32355822 | 153,414,160 | 2.3E-06 | Inact | *Ajap1* | 13,856 |  |  |  |
| 5 | rs6393330 | 138,877,875 | 3.0E-06 | % Act Press | *Gna12* | 1,916,545 |  | N |  |
| 11 | rs26988786 | 54,860,681 | 2.9E-08 | Inact | *Hint1* | 7,761 |  |  |  |
| 11 | rs26942304 | 60,457,090 | 2.8E-06 | Act Press | *Drg2* | 4,582 |  |  | N, mF |
| 11 | rs26986383 | 60,484,778 | 1.1E-07 | Infus | *Drg2* | -23,106 |  |  | N, mF |
| 11 | rs26984580 | 62,605,569 | 3.2E-06 | Act Press | *Trpv2* | -18,069 | N | N | N, mF |
| 11 | rs26984580 | 62,605,569 | 8.3E-09 | Infus | *Trpv2* | -18,069 | N | N | N, mF |
| 11 | rs28241639 | 81,647,142 | 3.4E-06 | % Act Press | *Asic2* | -222,829 |  |  | mF |
| 12 | rs33619289 | 114,955,853 | 2.4E-07 | Act Press | *Vipr2* | 1,156,141 |  |  |  |
| 13 | rs29602391 | 21,073,361 | 3.2E-06 | % Act Press | *Trim27* | 113,723 |  |  |  |
| 14 | rs48220977 | 56,380,012 | 3.3E-06 | Inact | *Rnf17* | 83,794 |  |  | mF |
| 17 | rs33126598 | 4,133,471 | 1.7E-06 | Inact | *Cldn20* | -600,587 |  |  | N |

<sup>1</sup> In the absence of other supporting evidence, candidate genes were nominated based on a combination of proximity and biological plausibility.
<sup>2</sup> Negative distance, gene centromeric to marker; positive, telomeric.
<sup>3</sup> FUSION in cocaine exposed mice; N, NAc; mF, mFC.
<sup>4</sup> FOCUS in cocaine exposed mice.
<sup>5</sup> *Cis* eQTLs in cocaine exposed mice.

Of the 15 unique loci, 8 had evidence from *cis* eQTLs or TWAS supporting the nomination of a nearby candidate gene (Table 1). Candidate genes for these loci included *A630001G21Rik, Plch1, Gna12, Drg2*, *Trpv2, Asic2, Rnf17* and *Cldn20*. For the remaining 7 loci, candidate genes were nominated based on proximity to the behavioral locus and biological plausibility. As expected, loci showed allelic differences in behavioral endpoints over the time course of the study (Supplementary Figure S11). Quantile-quantile (QQ) plots showed deviations from normality, reflecting the complex genetic structure of the HMDP and the longitudinal nature of the phenotypes (Supplementary Figure S12) (Tyler et al., 2021).

### RNA-Seq

To better discern pathways for cocaine self-administration, RNA-Seq was performed on NAc and mFC from 41 of the cocaine- and saline-exposed strains from the HMDP. The NAc and mFC were chosen because of the enhanced dopaminergic signaling in these brain regions caused by cocaine, which is responsible for much of the drug’s action (Fernàndez-Castillo et al., 2022; Pierce et al., 2018). A total of 72.6 ± 1.1 M paired-end reads were obtained per strain in each brain region for cocaine and 73.0 ± 1.1 M for saline (Bagley et al., 2022).

Principal components analyses were performed using transcript abundance, splicing (percent spliced in, psi or ψ) and RNA editing (percent edited, phi or ϕ) (Supplementary Figure S13). Samples showed strong separation due to region, some separation due to batch, but little or no separation based on sex or infusate. A total of four samples showed possible misattribution based on region, corresponding to an error rate of 1%. This error rate is comparable to, or better than, other genome-scale studies (Broman et al., 2015; Westra et al., 2011). To avoid over-fitting, we elected not to correct the putatively misassigned samples.

### Gene regulation

Changes in transcript and isoform abundance may be caused directly by the infusate or indirectly influenced by the various genetic backgrounds. Our study is nearly balanced with respect to infusate and mouse strain, so genetic background is unlikely to be an appreciable confound. Nevertheless, to ensure that we identified expression changes independent of genetic background, we used a linear mixed model implemented in `lme4qtl` to correct for population structure. Further, the model was omnibus in nature, with fixed effects of brain region, sex, infusate, sex/infusate interaction and batch. This approach allowed a unified approach to the identification of transcript abundance changes caused by the relevant biological effects. For splicing and RNA editing, a fixed effect of transcript expression level was added to the model. This addition enabled us to identify significant links between splicing or RNA editing and transcript abundance.

### Genes regulated by cocaine

A total of 5,111 transcripts were significantly regulated by cocaine, including *Per2*, *Fam107a*, *Eif5* and *Ankrd28* (false discovery rate, FDR < 0.05) (Figure 2A, Supplementary Figure S14). *Per2* is a core circadian rhythm gene known to be regulated by cocaine that, in turn, alters the effects of cocaine on circadian phase shifts (Prosser et al., 2014). Gene ontology (GO) analysis of transcripts regulated by cocaine using the biological process term showed 465 significant functional categories (FDR < 0.05) including nitrogen compound and organic substance metabolic process (Supplementary Figure S15).

**Figure 2.**
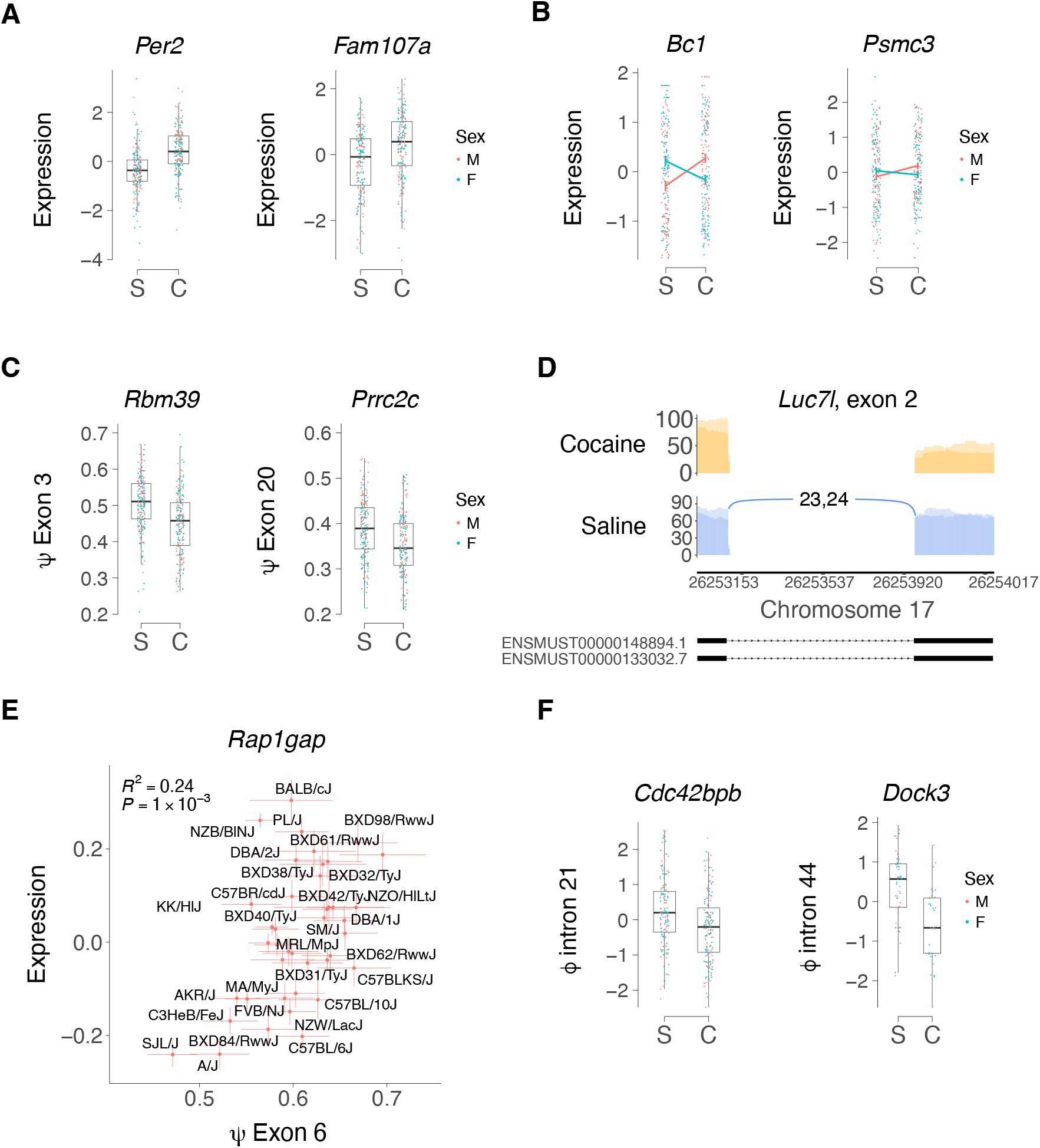
Regulation of gene expression by cocaine. (**A**) Transcript abundance. Per2, FDR < 2.2 × 10^−16^; Fam107a, FDR < 2.2 × 10^−16^. S, saline; C, cocaine. M, male; F, female. (**B**) Sex/infusate interactions for transcript abundance. Bc1, FDR = 3.8 × 10^−12^; Psmc3, FDR = 3.5 × 10^−3^. Means ± s.e.m. (**C**) Splicing. Rbm39, exon 3, FDR = 2.2 × 10^−12^; Prrc2c, exon 3, FDR = 4.0 × 10^−6^. (**D**) Sashimi plot (Garrido-Martín et al., 2018) of cocaine-regulated splicing of Luc7l, exon 2. FDR = 7.6 × 10^−3^. (**E**) Alternate splicing of Rap1gap, exon 6, affects transcript abundance. Strain means ± s.e.m. R^2^ and P values are strain averaged results; linear mixed model FDR < 2.2 × 10^−16^. (**F**) RNA editing. Cdc42bpb, editing site Chromosome 12, 111,309,987 bp in Alu element B1_Mus2, intron 21, FDR = 0.04; Dock3, editing site Chromosome 9, 106,905,884 bp in Alu element B1_Mur1, intron 44/49, FDR = 0.04.

Confirming our results, there was significant overlap between the regulated transcripts found in our investigation and a recent study that used RNA-Seq to examine the NAc of C57BL/6J mice exposed to cocaine IVSA (odds ratio = 1.7, *P* = 6.2 × 10^−5^, Fisher’s Exact Test) (Walker et al., 2018). We found a total of 48 transcripts with significant sex/infusate interactions (FDR < 0.05), including *Bc1*, *Crebzf*, *Taok1* and *Psmc3* (Figure 2B, Supplementary Figure S14). *Bc1* is a non-coding RNA gene that is transported to dendrites where it is involved in translational regulation (Robeck et al., 2016).

A total of 31 exons showed significant differential splicing for cocaine compared to saline (FDR < 0.05) (Figure 2C,D, Supplementary Figure S16). Examples included *Rbm39* and *Luc7I*, which themselves both regulate splicing (Daniels et al., 2021; Xu et al., 2022). Reminiscent of *Per2*, *Rbm39* shows a circadian rhythm-based regulation of splicing (El-Athman et al., 2019). A total of 860 differential splicing events resulted in altered transcript abundance levels, most likely by affecting RNA stability (FDR < 0.05) (Figure 2E, Supplementary Figure S17) (Baralle and Giudice, 2017). GO analysis of spliceforms regulated by cocaine revealed significant enrichment in terms related to RNA splicing (Supplementary Figure S18).

Only two genes showed significant changes in RNA editing levels as a result of cocaine, *Cdc42bpb* and *Dock3* (FDR < 0.05) (Figure 2F, Supplementary Figure S19). Both editing sites are in intronic *Alu* elements. GO analysis of genes in which RNA editing was nominally regulated by cocaine or cocaine/sex interactions (*P* < 0.05) showed significant enrichment of a number of categories including organelle organization, metabolic processes, including nitrogen compound metabolic process, and RNA processing (Supplementary Figure S20).

### Genes regulated by brain region and sex

Significant regulation of gene expression due to brain region or sex was identified for transcript abundance (brain region, 19,297 transcripts; sex, 822; FDR < 0.05), spliceforms (brain region, 1,275 spliceforms; sex, 6) and RNA editing (brain region, 299 editing sites; sex, 0) (Supplementary Figures S14, S16 and S19). Brain region had a larger number of regulated gene expression events than either cocaine or sex. Significant GO enrichments were found for brain region and sex, including cellular and primary metabolic process for brain region, and chromosome organization and histone modification for sex (Supplementary Figures S15, S18 and S20).

Of the 299 editing sites significantly regulated by brain region, 13 resulted in non-synonymous coding region changes, including *Cadps*, *Tmem63b*, *Unc80* and *Cyfip2* (Supplementary Figure S19) (Cuddleston et al., 2022; Shumate et al., 2021; Tariq et al., 2013; Wu et al., 2020).

### Expression QTLs

*Cis* and *trans* QTLs were identified for transcript abundance (expression QTLs, or eQTLs), splicing (sQTLs or ψQTLs) and RNA editing (edit QTLs or ϕQTLs) using `FaST-LMM` (Supplementary Figures S21-S23). The number of *cis* eQTLs averaged over the two brain regions and infusates was 4,844 ± 159 (Figure 3A, Supplementary Figure S21). The distance between the *cis* eQTLs and their corresponding gene was 0.63 Mb ± 0.005 Mb, averaged across brain regions and infusates, consistent with the known linkage disequilibrium structure of the HMDP (Figure 3B, Supplementary Figure S24) (Bennett et al., 2010; Ghazalpour et al., 2012; Lusis et al., 2016).

**Figure 3.**
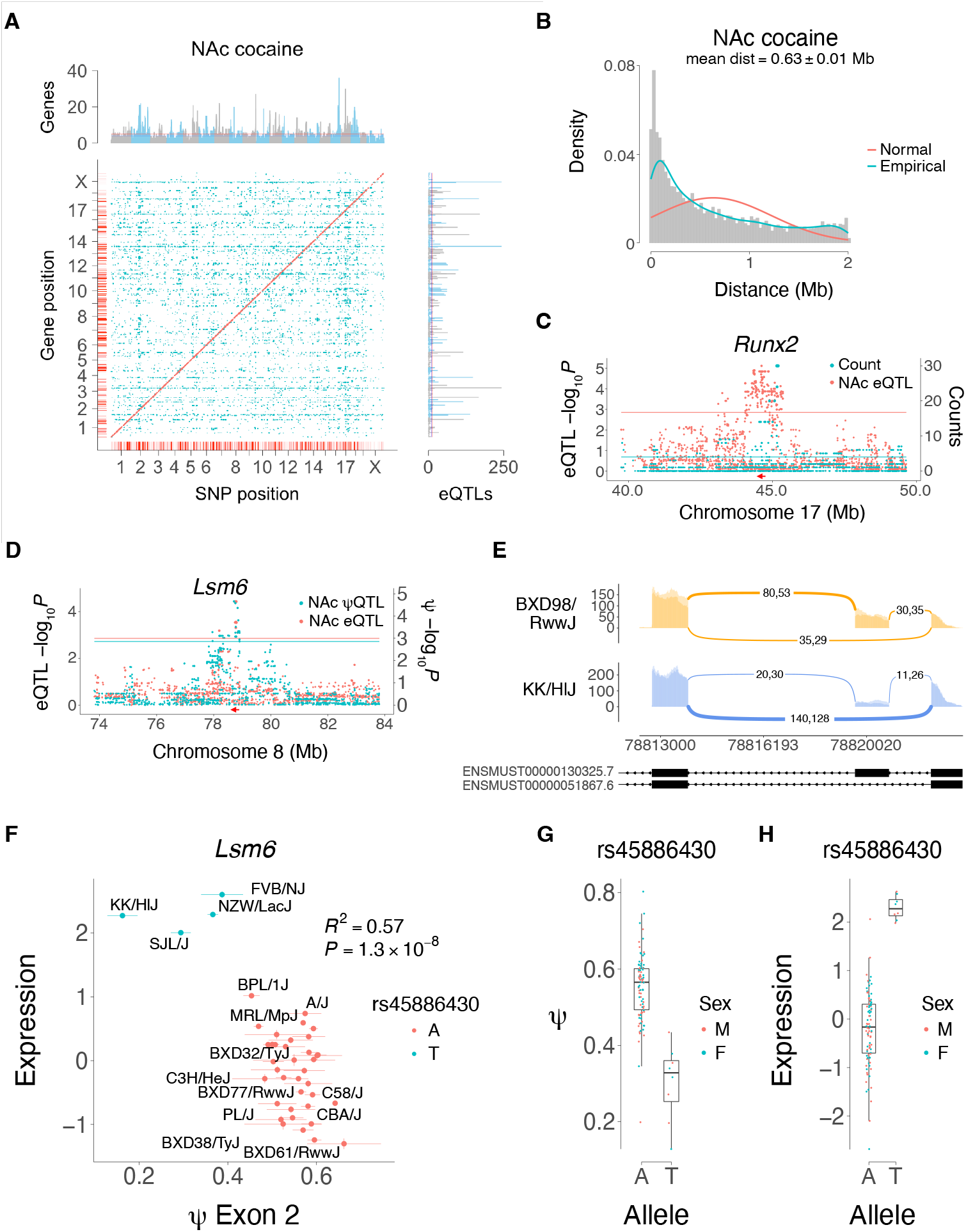
Genetic regulation of gene expression in NAc from cocaine-exposed mice. (**A**) Cis (red) and trans (blue) eQTLs. Marginal graphs show SNPs regulating many genes (horizontal graph, eQTL hotspots) and genes regulated by many eQTLs (vertical graph). Blue lines, FDR < 0.05, red lines, FDR < 0.01 (Poisson). Distance between cis eQTLs and the corresponding genes. (**C**) Co-aligned eQTL hotspot and Runx2 cis eQTL. Red arrow, location of Runx2. Blue horizontal line, eQTL hotspot significance threshold, FDR < 0.05. Red horizontal line, cis eQTL significance threshold. (**D**) Coincident cis ψQTL for exon 2 and eQTL for Lsm6. Peak marker rs45886430 for both QTLs. Blue and red horizontal lines, respective significance thresholds. (**E**) Sashimi plot for exon 2 of Lsm6 in BXD98/RwwJ and KK/HIJ, with A or T allele of rs45886430, respectively. (**F**) Allele A of rs45886430, the SNP most strongly correlated with both alternate splicing of exon 2 and expression of Lsm6, is associated with higher ψ for exon 2 of Lsm6 and lower expression. Means ± s.e.m. for each strain. (**G**) Allele effect of rs45886430 on ψ of Lsm6 exon 2. Individual samples shown. (**H**) Allele effect of rs45886430 on Lsm6 expression.

We identified hotspots for transcript abundance, in which a locus regulates many genes (Hasin-Brumshtein et al., 2016; Park et al., 2011a). A total of 9 hotspots regulating ≥ 20 genes were present in NAc cocaine samples (FDR < 2.2 × 10^−16^), and 10 hotspots in mFC cocaine samples (FDR < 2.2 × 10^−16^). We sought candidate genes for hotspots by looking for co-aligned *cis* eQTLs. One NAc cocaine hotspot was coincident with a *cis* eQTL for the transcription factor *Runx2* (Figure 3C).

### Splicing QTLs

A total of 1,426 ± 20 *cis* splicing QTLs (ψQTLs) were detected, averaged over the two brain regions and infusates (Supplementary Figure S22). Spliceforms regulated by genetic variants can affect transcript abundance as a result of changes in mRNA stability (Mockenhaupt and Makeyev, 2015; Titus et al., 2021). To evaluate the prevalence of this phenomenon, we examined whether there was a statistically significant enrichment in coincident *cis* eQTLs and ψQTLs. There were 360 coincident *cis* eQTLs and ψQTLs in NAc from cocaine treated mice (odds ratio, OR, = 2.8, *P* < 2.2 × 10^−16^, Fisher’s exact test), while cocaine treated mFC had 419 (OR = 2.6, *P* < 2.2 × 10^−16^). An example of a coincident *cis* eQTL and ψQTL for *Lsm6* in NAc of cocaine exposed mice is shown in Figures 3D-H. *Lsm6* is involved in pre-mRNA splicing (Wu et al., 2012).

### Editing QTLs

RNA editing results in sequence changes that can affect transcript stability and abundance and can also alter coding sequence (Brümmer et al., 2017). We identified 272 ± 11 *cis*-acting loci that affect RNA editing efficiency (ϕQTLs), averaged over the two brain regions and infusates (Supplementary Figure S23) (Li et al., 2022). Because RNA editing occurs at single nucleotides, confident quantitation of these events is more demanding than transcript or spliceform abundance. The ascertainment rate for all editing events was 37 ± 0.4% of RNA-Seq samples, averaged across infusates and brain regions. The decreased power resulting from the < 100% detection rate means that the ϕQTLs should be treated with some caution.

To evaluate how often genetically driven variations in RNA editing can affect transcript abundance, we tested for statistically significant enrichment in coincident *cis* eQTLs and ϕQTLs. There were 51 coincident *cis* eQTLs and ϕQTLs in NAc from cocaine treated mice (odds ratio, OR, = 2.0, *P* = 4.2 × 10^−5^, Fisher’s exact test), while cocaine treated mFC had 46 (OR = 1.6, *P* = 5.3 × 10^−3^). If *cis* ϕQTLs regulate editing events that in turn alter transcript stability and give rise to *cis* eQTLs, coincident *cis* ϕQTLs and *cis* eQTLs should be enriched in editing sites that appear in the final transcript rather than intronic or intergenic regions. This prediction was fulfilled. We found significant enrichment of editing sites affecting 5′ untranslated, 3′ untranslated and exonic coding regions in the coincident *cis* ϕQTLs and eQTLs in both cocaine exposed NAc (odds ratio = 1.9, *P* = 9.2 × 10^−3^, Fisher’s Exact Test) and cocaine exposed mFC (odds ratio = 2.9, *P* = 2.4 × 10^−5^, Fisher’s Exact Test).

A *cis* ϕQTL that regulates editing of a site in a B1_Mm *Alu* element in the 3′ untranslated region of *Samd8* (Chromosome 14, 21,797,711 bp) and that aligns with a *cis* eQTL in NAc of cocaine exposed mice is shown in Figure S25.

### Integrating RNA-Seq and behavioral loci

To identify candidate genes for cocaine self-administration, we sought eQTLs that aligned with the behavioral loci. Of the 15 unique IVSA loci, 9 aligned with either cocaine exposed NAc or mFC *cis* eQTLs (Table 1). The two most significant loci for cocaine infusions mapped to Chromosome 11 at 60,484,778 bp and 62,605,569 bp (Figures 1E-H and 4). Plausible candidate genes for the two loci were *Drg2* and *Trpv2*, respectively, each of which were supported by significant *cis* eQTLs in both NAc and mFC (Figure 4). Higher expression of both *Drg2* and *Trpv2* was associated with lower cocaine infusions. There was, however, linkage disequilibrium between the *Drg2* and *Trpv2* loci (*D*′ = 0.91, *R*^2^ = 0.67, *P* < 2.2 × 10^−16^), making it hard to disentangle their relative contributions. Other genes supported by co-aligned *cis* eQTLs in cocaine IVSA loci included *A630001G21Rik*, *Plch1*, *Asic2*, *Rnf17* and *Cldn20* (Table 1).

**Figure 4.**
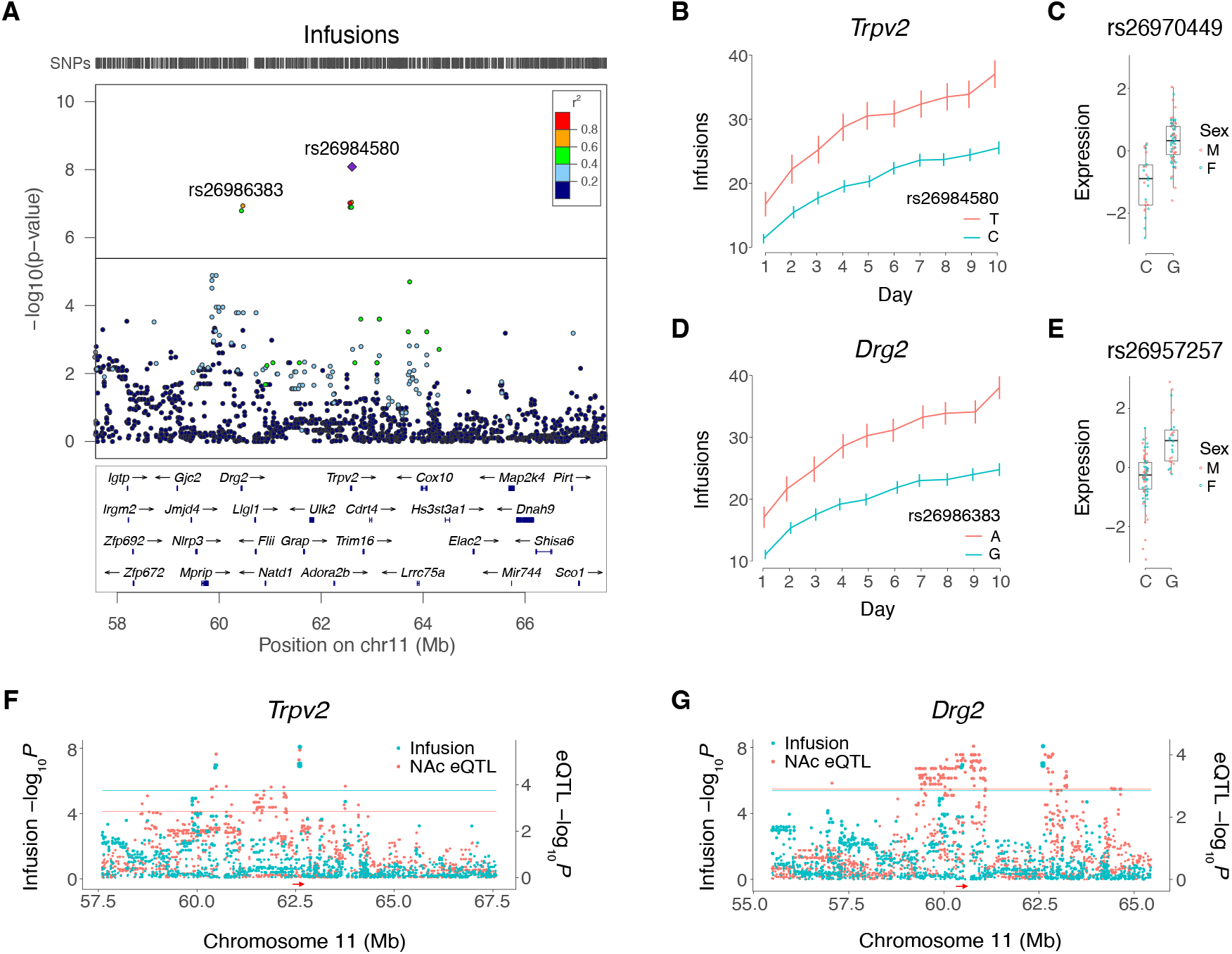
Trpv2 and Drg2 are close to infusion QTLs and have cis eQTLs. (**A**) Locuszoom plot for cocaine IVSA infusions, showing loci harboring Drg2 and Trpv2 (Pruim et al., 2010). R^2^ values convey linkage disequilibrium. (**B**) Infusion time course for peak SNP of Trpv2 locus, rs26984580. Means ± s.e.m. Expression of Trpv2 for peak cis eQTL SNP, rs26970449, in cocaine-exposed NAc. SNPs rs26984580 and rs26970449 are in linkage disequilibrium (D′ = 1, R^2^ = 0.71, P < 2.2 × 10^−16^). (**D**) Infusion time course for peak SNP of Drg2 locus, rs26986383. (**E**) Expression of Drg2 for peak cis eQTL SNP, rs26957257, in cocaine-exposed NAc. SNPs rs26986383 and rs26957257are in linkage disequilibrium (D′ = 1, R^2^ = 0.07, P = 1.8 × 10^−9^). (**F**) Coincident loci for infusions and Trpv2 cis eQTL in cocaine-exposed NAc. (**G**) Coincident loci for infusions and Drg2 cis eQTL in cocaine-exposed NAc.

### Transcriptome-wide association studies

We used transcriptome-wide association studies (TWASs) to further evaluate genetic pathways for cocaine IVSA. The TWAS approach identifies significant *cis* eQTLs and nominates the corresponding gene if there is significant correlation of its expression with a trait. Because TWAS analyzes pathways at the gene rather than the marker level, there is decreased multiple hypothesis correction and the approach provides increased statistical power. We used FUSION and FOCUS software to perform TWAS, with FOCUS providing additional fine mapping (Gusev et al., 2016; Mancuso et al., 2019).

Consistent with its increased statistical power, TWAS identified 20 significant genes for cocaine IVSA, of which 17 were supplementary to the genetic loci (Table 1, Figure 5, Supplementary Tables S2-S3, Supplementary Figures S26-S27). Of the 20 TWAS significant genes, 12 were significant using FUSION (*Slco5a1, Cpxm1, Gm14057, Arhgef26, Tprkb, Slc18b1, Mgat4b, Hnrnpab, Gdi2, Trat1, Dubr* and *Gm10232*), 5 were significant using FOCUS (*Slc4a11, Gna12, 9930111J21Rik2, Gm12216* and *Mief2*) and 3 (*A630001G21Rik*, *G3bp1* and *Trpv2*) were common to both.

**Figure 5.**
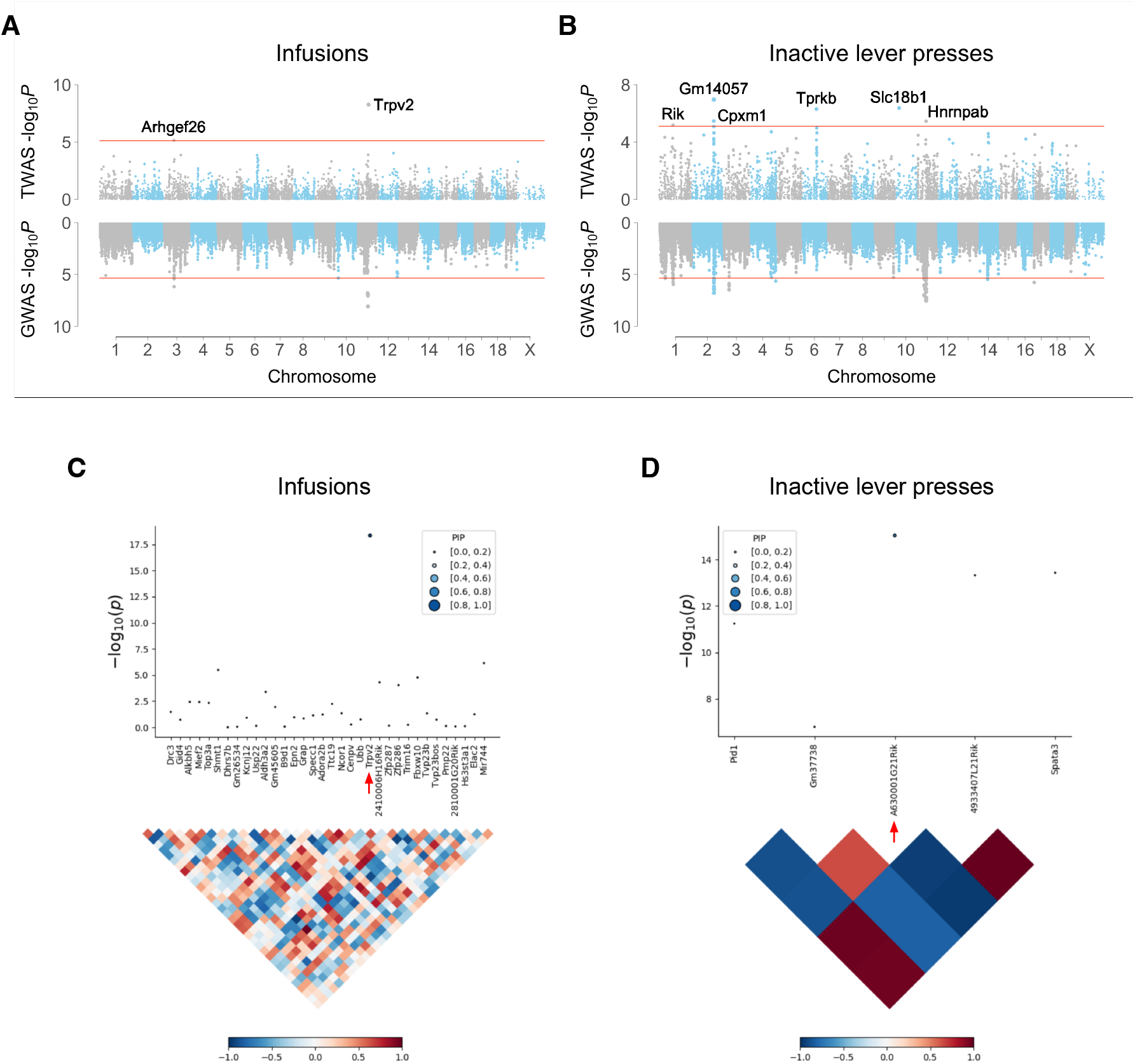
TWAS for cocaine infusions and inactive lever presses using NAc RNA-Seq data from cocaine-exposed mice. (**A**) FUSION TWAS of infusions highlights Trpv2. (**B**) FUSION TWAS of inactive lever presses. Rik, A630001G21Rik. (**C**) FOCUS of Trpv2 locus (red arrow) for infusions. Linkage disequilibrium map shown underneath. (**D**) FOCUS of A630001G21Rik locus for inactive lever presses.

In some cases, TWAS suggested different genes than those nominated on the basis of proximity and biological plausibility (Table 1). For example, *G3bp1* was significant for cocaine inactive lever presses using both FUSION (*P* = 1.83 × 10^−6^) and FOCUS (pip = 0.91) on RNA-Seq data from cocaine-exposed mFC. However, *G3bp1* was 626,581 bp from the nearest behavioral QTL. In contrast, the nominated candidate gene for this QTL, *Hint1*, was 7,761 bp from the locus. Although *Hint1* had no significant *cis* eQTLs, we gave this gene precedence because of its proximity to the behavioral QTL and its known role in addiction (Barbier et al., 2007; Liu et al., 2017). Further work is required to distinguish which of the two genes (or both) is relevant to cocaine IVSA. A similar situation exists for *Pdyn* and *Cpxm1*, and *Drg2* and *Mief2*. Regardless, the TWAS genes for cocaine IVSA are useful entry points for new studies of drug addiction.

TWAS provided support for 3 genes in the 15 non-redundant cocaine IVSA loci (Table 1); *A630001G21Rik*, *Gna12* and *Trpv2*. FUSION and FOCUS provided strong evidence for *Trpv2* in both cocaine infusions and active lever presses using cocaine-exposed NAc but not cocaine-exposed PFC data. However, *Drg2* was not supported by TWAS, despite significant *cis* eQTLs in cocaine-exposed NAc and mFC. Additional strain transcriptomes may provide enough power to support a role for *Drg2* in cocaine IVSA using TWAS. Alternatively, *Drg2* may exert its effects through amino acid variations rather than expression, although no such variants are currently known among the 37 sequenced inbred mouse strains (Lilue et al., 2018; Timmermans et al., 2017). Genome sequencing of further strains in the HMDP may reveal *Drg2* protein-altering variants.

### eCAVIAR

To further dissect the contributions of *Trpv2* and *Drg2* to cocaine IVSA, we used eCAVIAR. This software evaluates the posterior probability that the same SNP is causal for both GWAS and expression QTLs, while accounting for the uncertainty introduced by linkage disequilibrium (Hormozdiari et al., 2016). A colocalization posterior probability (CLPP) > 0.01 supports sharing of causal GWAS and expression QTL variants.

Consistent with the TWAS results, *Trpv2* had above-threshold CLPPs for infusions (NAc, rs26984580, CLPP = 0.13; mFC, rs26970449, CLPP = 0.03) and active lever presses (NAc, rs26984580, CLPP = 0.06; mFC, rs26970449, CLPP = 0.02) while *Drg2* did not. Also consistent with TWAS (Table 1), CLPPs for *Trpv2* were higher in cocaine-exposed NAc than mFC, suggesting NAc is the more relevant target tissue. The SNP with the highest CLPP for *Trpv2* (rs26984580) was 18 kb telomeric to *Trpv2* and located 180 bp telomeric to a proximal enhancer-like element and 182 bp centromeric to a promoter-like element in the 3′ untranslated region of the neighboring gene, *Lrrc75a*.

## Discussion

We integrated a longitudinal dataset of cocaine and saline IVSA with RNA-Seq to dissect genetic pathways for cocaine use in mice. Four behavioral endpoints were employed. Infusions, active lever presses and percent active lever presses evaluate how hard the animal is willing to work for delivery of cocaine or saline. Percent active lever presses control for overall locomotor activity by normalizing active lever presses to total lever presses. In contrast, inactive lever presses may measure either the aversive properties of the infusate or locomotor activity, whether intrinsic or modified by infusate.

Longitudinal GWAS increased the power to detect QTLs affecting cocaine IVSA. GWAS using individual days identified no genome-wide significant loci for any of the four behavioral endpoints, while GWAS using two day intervals identified two genome-wide significant loci for infusions (Bagley et al., 2022). The longitudinal GWAS identified a total of 15 unique loci using the four IVSA endpoints, of which three were identified for cocaine infusions. Further, the longitudinal GWAS QTLs had higher peak −log_10_*P* values than the GWAS using two day intervals.

Support for candidate genes in the longitudinal GWAS of cocaine IVSA was provided by the existence of coincident behavioral loci and *cis* eQTLs. TWAS gave further support for candidate genes in the GWAS, while suggesting additional candidates. Identification of *cis* ψQTLs and *cis* ϕQTLs that were coincident with *cis* eQTLs allowed documentation of numerous instances where alternate splice site selection or RNA editing altered transcript abundance, possibly by changing mRNA stability.

Of the 15 unique loci for cocaine IVSA identified by longitudinal GWAS, TWAS provided support for 3 candidate genes, while identifying a further 17 genes. *Trpv2* was close to the strongest GWAS locus for cocaine infusions as well as a locus for active presses. The role of *Trpv2* in cocaine IVSA was further supported by *cis* eQTLs in NAc and mFC, TWAS and eCAVIAR. The TWAS and eCAVIAR analyses suggested NAc was a more relevant target tissue than mFC.

*Trpv2* is a cation channel that is activated by cannabidiol and Δ9-tetrahydrocannabinol, possibly explaining the overlapping genetic risk factors for cocaine and cannabis use disorders (Kendler et al., 2007; Muller et al., 2019; Qin et al., 2008). In fact, the six known Trp cation channels are sometimes referred to as ionotropic cannabinoid receptors. Cannabidiol shows promise as a treatment for cocaine use disorder, consistent with our observation that higher expression of *Trpv2* is associated with lower cocaine IVSA (Calpe-López et al., 2019).

A second locus for infusions and active presses on Chromosome 11 was 2,120,791 bp centromeric to the *Trpv2* locus and was close to *Drg2*. Support for *Drg2* as a gene for cocaine IVSA was provided by *cis* eQTLs in NAc and mFC, but not by TWAS or eCAVIAR. Consistent with a potential role in cocaine addiction, *Drg2* knockout mice showed decreased dopamine release in the striatum (Lim et al., 2019). Knockout mice have also been created for *Trpv2* and show macrophage and mechanical nociception phenotypes (Entin-Meer et al., 2017; Katanosaka et al., 2018; Park et al., 2011b). Both the *Trpv2* and *Drg2* knockout mice would be attractive reagents to test the role of these genes in cocaine IVSA.

A locus for inactive lever presses on Chromosome 1 was close to *A630001G21Rik*, a likely nuclear body protein involved in transcription, and was supported by NAc and mFC *cis* eQTLs and also TWAS (Blake et al., 2021). Another locus for inactive lever presses was close to *Pdyn* on Chromosome 2. Although not supported by *cis* eQTLs or TWAS, *Pdyn* encodes prodynoprhin, which is proteolytically processed to produce κ opioid receptor ligands and modulates the response to cocaine (Abraham et al., 2022).

Our previous GWAS evaluating cocaine infusions for binned two day intervals identified a significant QTL on Chromosome 14 (56,388,089 bp) and on Chromosome 3 (51,706,218 bp). A suggestive QTL was also detected on chr 3 (36,771,265 bp). The Chromosome 14 QTL is 8,077 bp telomeric to a QTL for inactive lever presses obtained using the longitudinal model. *Rnf17* was close to this QTL and was supported by a *cis* eQTL in mFC. The suggestive QTL for infusions on Chromosome 3 using two day intervals is 1,406,935 bp centromeric to a significant locus for inactive lever presses obtained using the longitudinal model that has *Spry1* as the nominated candidate gene.

Candidate genes lacking additional supporting evidence in the current study are speculative. For example, the peak SNP for the locus on chromosome 12 regulating the number of active presses (rs33619289) is actually located in the middle of the immunoglobulin heavy chain locus. The nominated candidate gene, *Vipr2*, is located 1,156,141 bp telomeric to the SNP. Copy number increases in *Vipr2* are associated with schizophrenia and alter dopaminergic neurotransmission in engineered mice (Tian et al., 2019). Another gene, *Ptprn2*, is 1,925,431bp telomeric to the peak SNP and is genome-wide significant in human GWAS for cognitive performance, risk taking and smoking initiation (Buniello et al., 2019). Knockout mice showed that absence of *Ptprn2* decreased brain dopamine, norepinephrine and serotonin concentrations (Nishimura et al., 2009).

Interestingly, *VIPR2* and *PTPRN2* were genome-wide significant in a linkage analysis of comorbid cocaine dependence and major depressive episode in humans (Yang et al., 2011). In our study, *Vipr2* had no *cis* eQTLs, while *Ptprn2* had significant *cis* eQTLs in both cocaine-exposed NAc and mFC. However, we nominated *Vipr2* rather than *Ptprn2* largely because *Vipr2* is substantially closer to the peak behavior SNP. Although both genes are biologically plausible and within the credible distance for linkage disequilibrium, their candidacy should be treated with caution.

Among the 17 genes identified by TWAS supplementary to GWAS, *Slc18b1* was significant for inactive lever presses using FUSION analysis of both cocaine-exposed NAc and mFC RNA-Seq data. *Slc18b1* is a serotonin transporter, consistent with the role of cocaine in inhibiting the reuptake of this neurotransmitter (Moriyama et al., 2020). Other TWAS significant genes for cocaine IVSA had relevant phenotypes in human GWASs. *Cpxm1* is genome-wide significant for cognitive and executive function; *Arhgef26* for body mass index, sulcal depth and smoking initiation; *G3bp1* for cortical thickness; and *Dubr* for worry (Buniello et al., 2019).

## Acknowledgements

Funding was provided from the National Institute on Drug Abuse, U01 DA041602, P50 DA039841; the National Institute on Alcohol Abuse and Alcoholism, T32 AA025606; and the National Institute of Mental Health, R01 MH123177. We thank the UCLA Semel Institute Neurosciences Genomics Core for sequencing. This work used computational and storage services associated with the Hoffman2 Shared Cluster provided by the UCLA Institute for Digital Research and Education Research Technology Group.

