## Supplementary Material for "Genetic pathways regulating the longitudinal acquisition of cocaine self-administration in inbred and recombinant inbred mice"

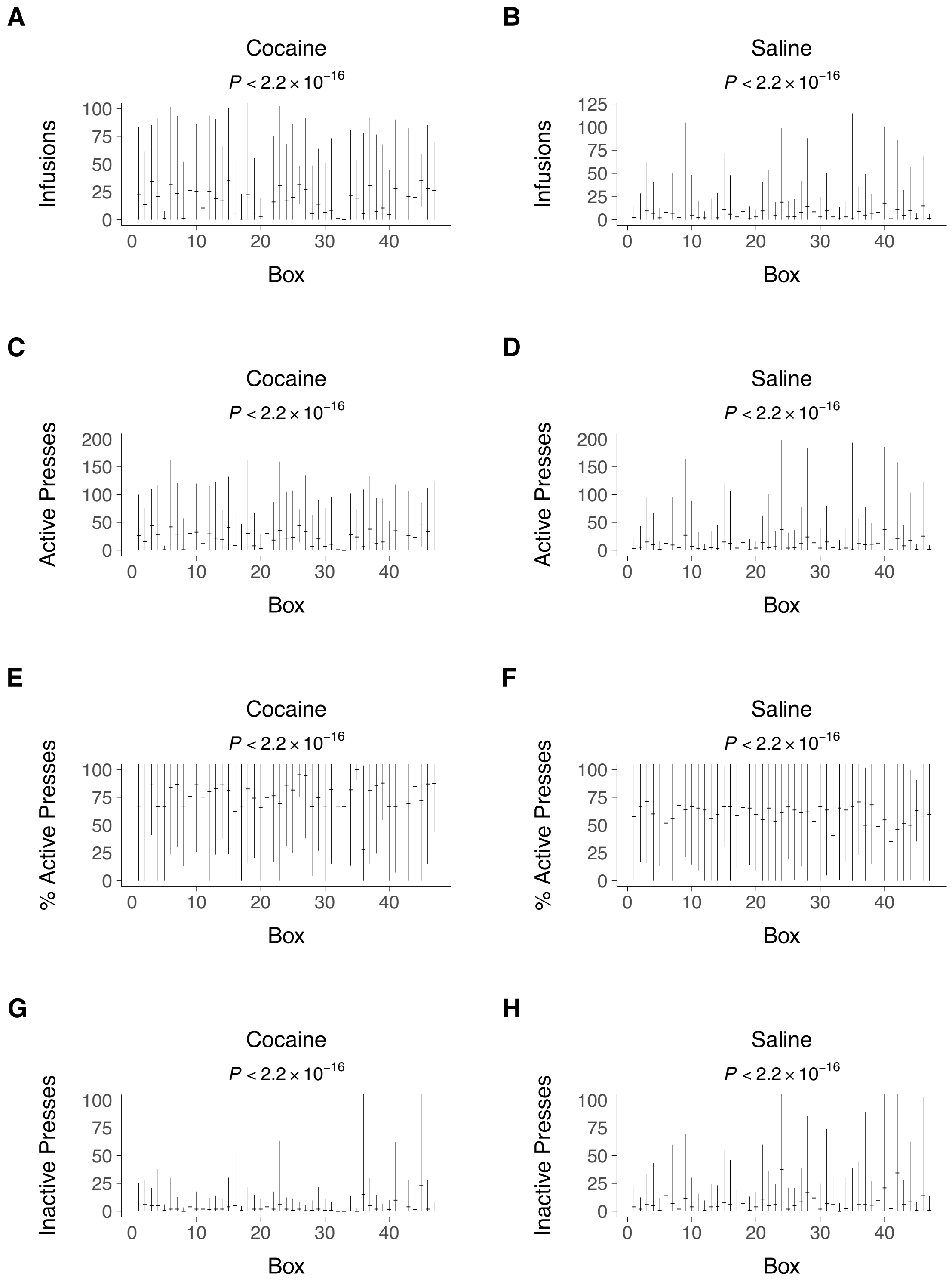


***Supplementary Figure S1.*** *Effects of self-administration chamber on cocaine and saline intravenous self-administration (IVSA). The horizontal bars correspond to the median. The upper limits of the whiskers corresponds to the median plus 1.75 × the interquartile range, and the lower limits to the median minus 1.75 × the interquartile range.*

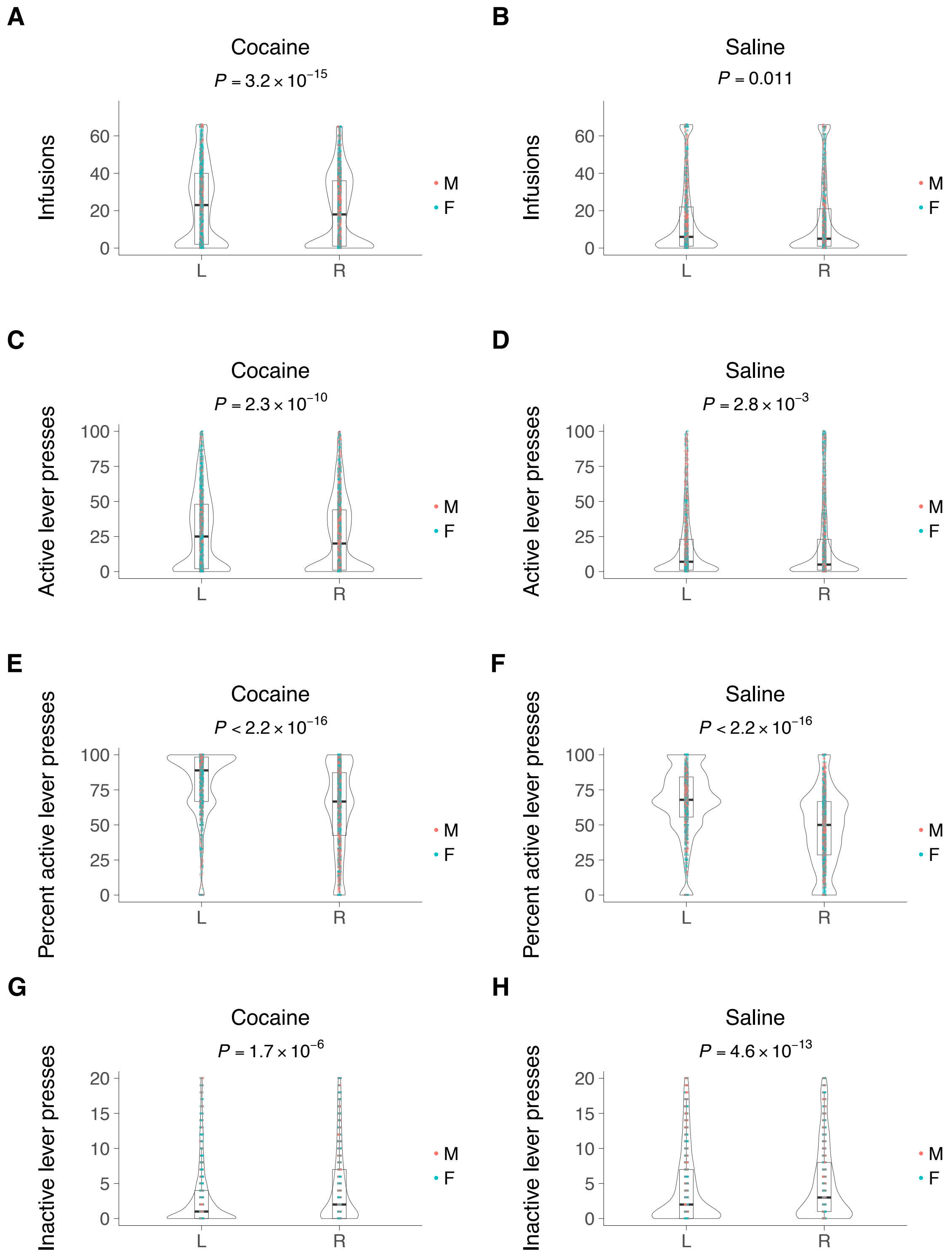


***Supplementary Figure S2.*** *Lever preferences. L, left; R, right. M, male; F, female.*

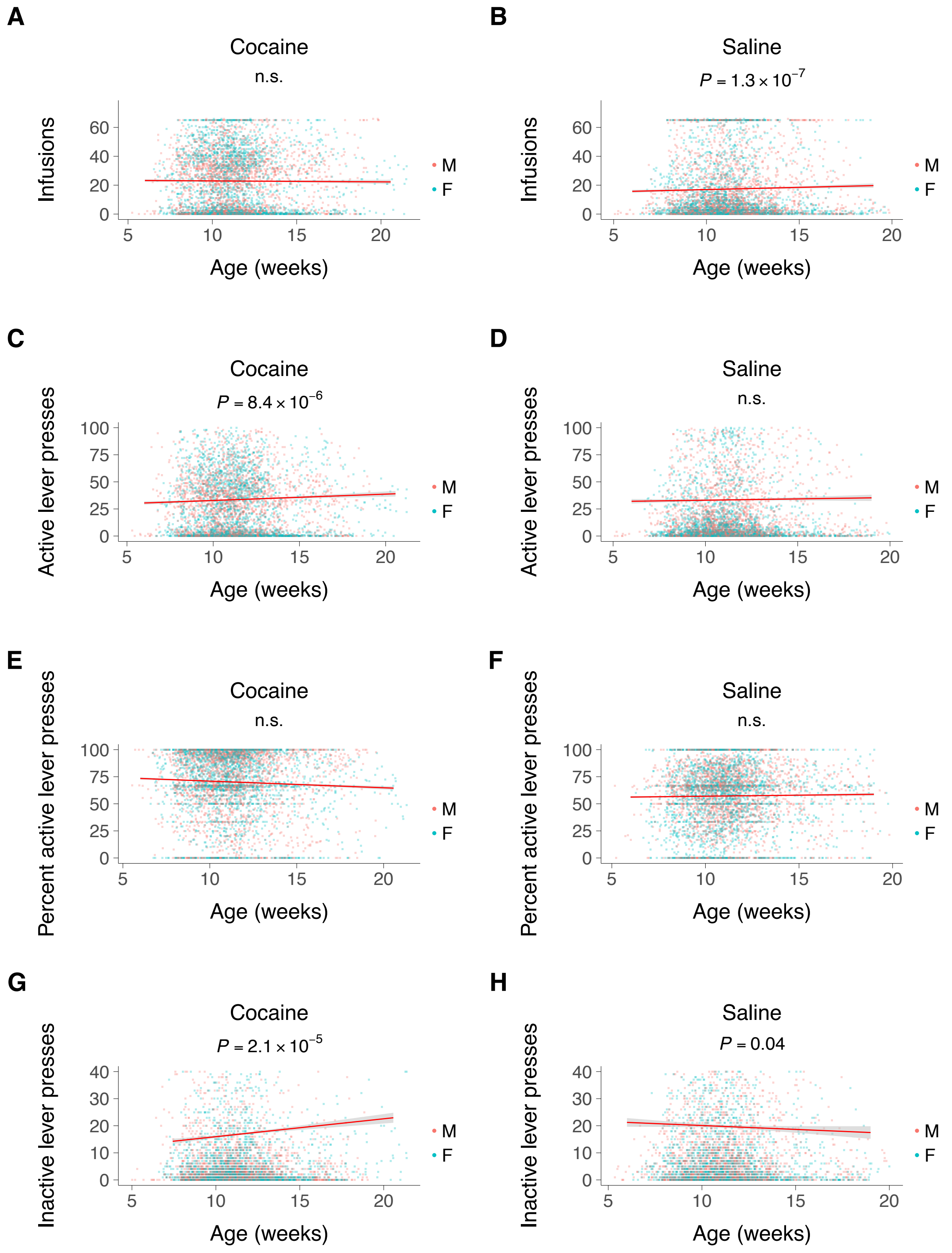


***Supplementary Figure S3.*** *Effects of age. M, male; F, female. Best fit with 95% confidence interval shown.*

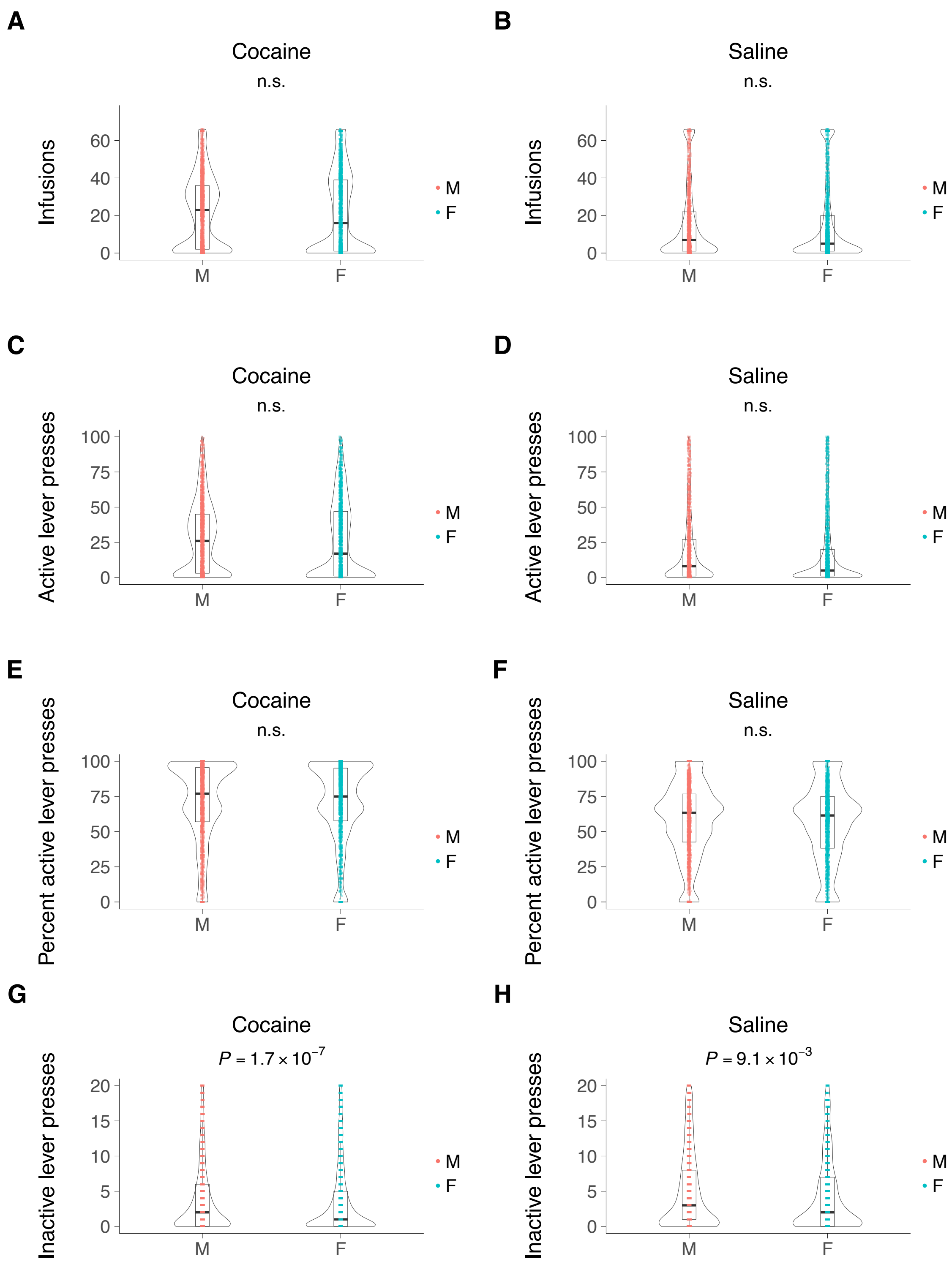


***Supplementary Figure S4.*** *Effects of sex. M, male; F, female.*

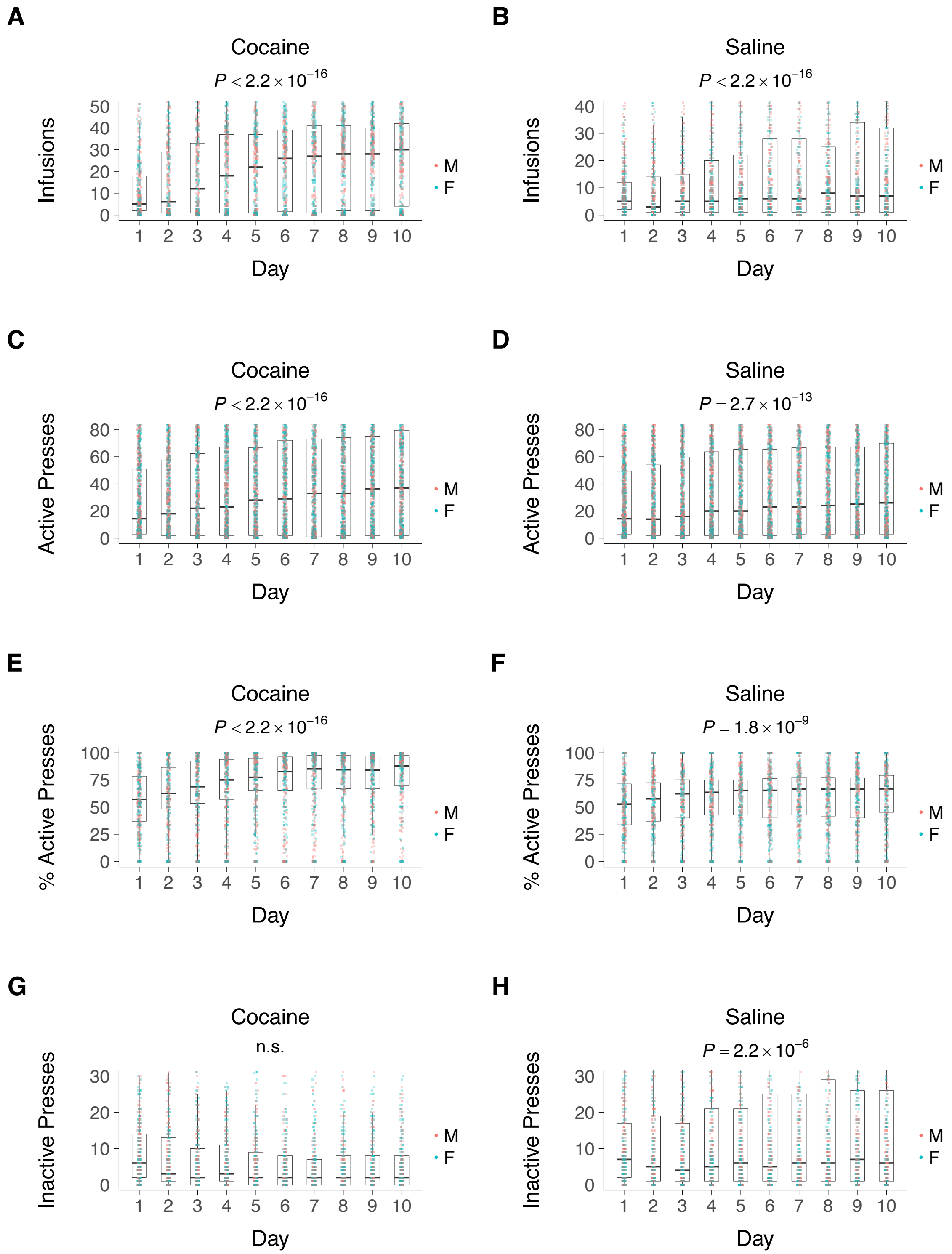


***Supplementary Figure S5.*** *Box plots showing self-administration of cocaine or saline. P values represent significance values for time. M, male; F, female.*

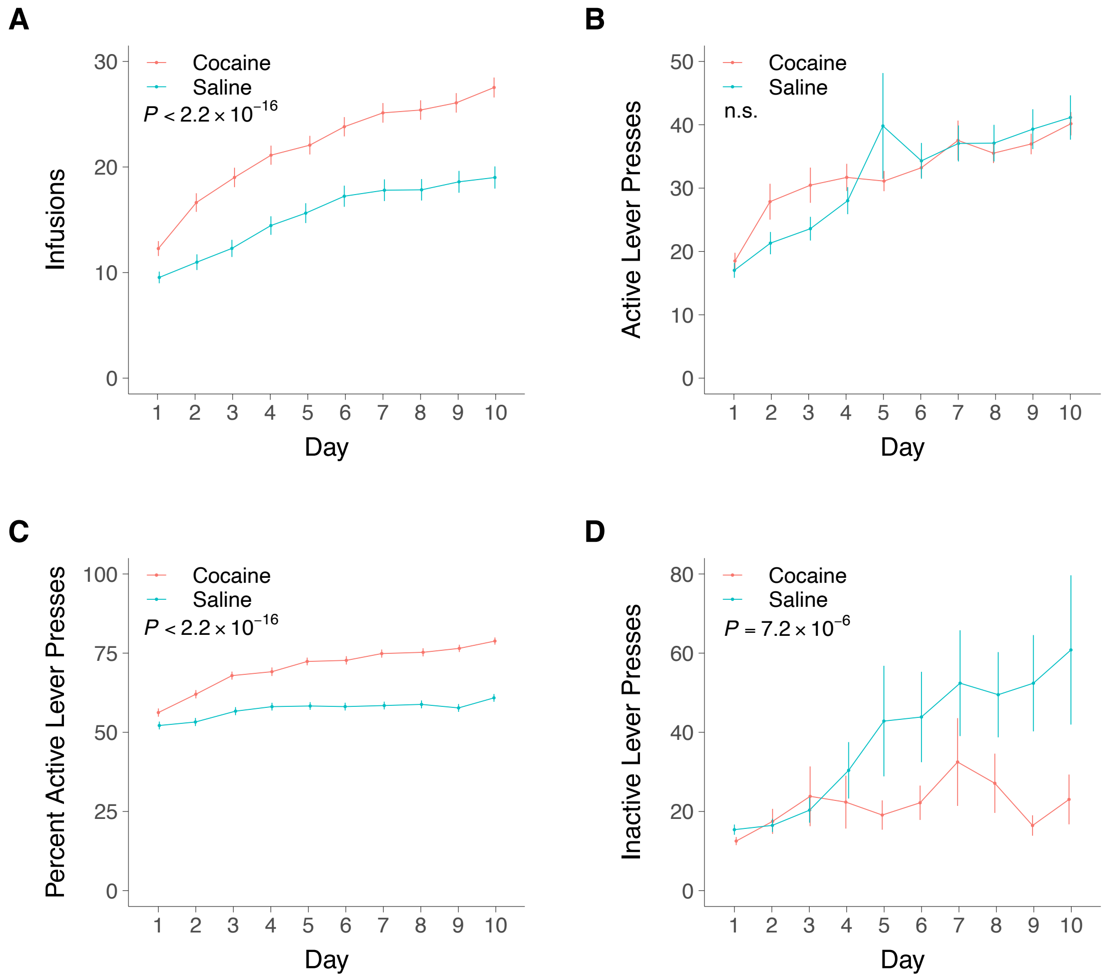


***Supplementary Figure S6.*** *Cocaine vs saline self-administration. Means ± s.e.m. P values represent comparisons of cocaine to saline.*

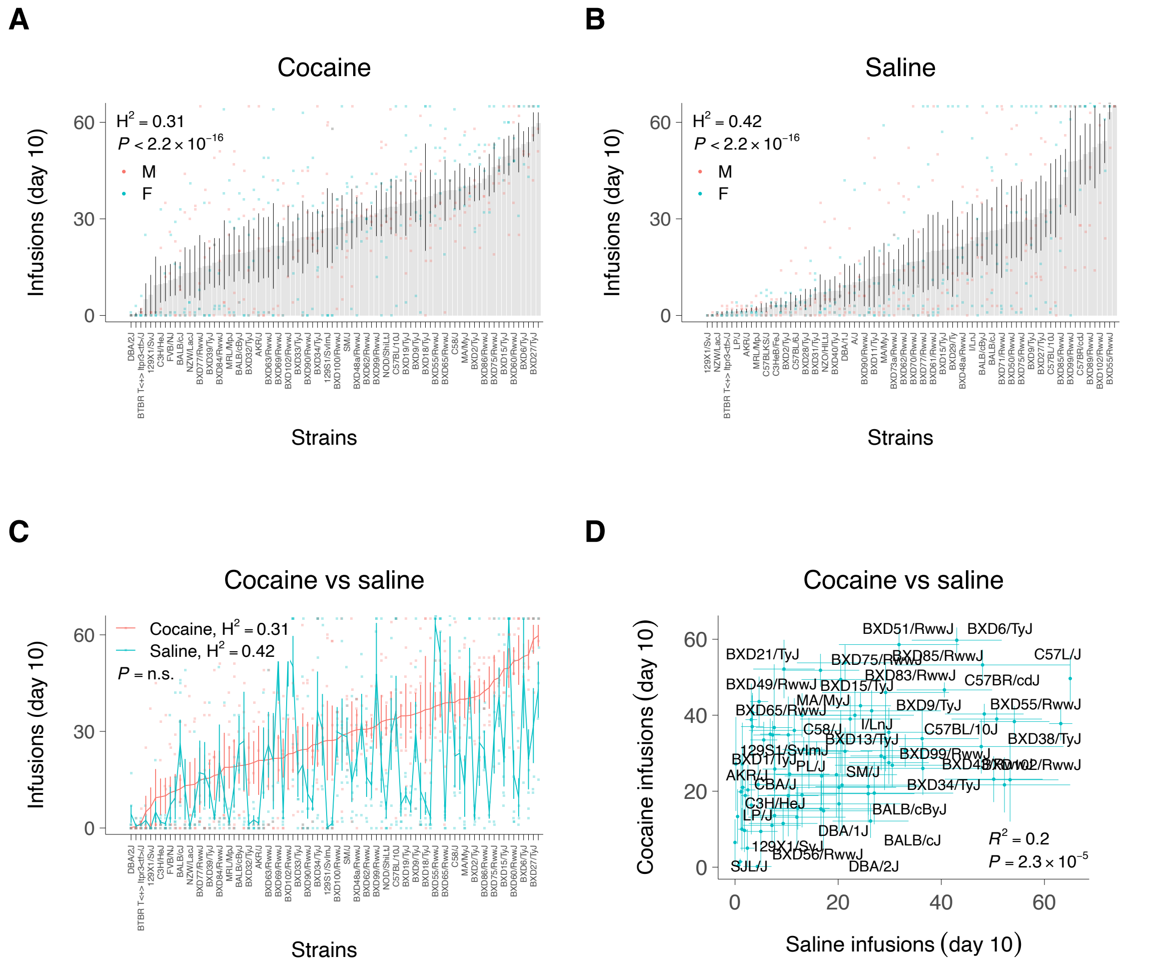


***Supplementary Figure S7.*** *Broad sense heritability ($H^2$) of self-administered infusions for cocaine and saline on day 10. (**A**) Strains ranked by cocaine infusions. P is significance of $H^2$. All strains shown, but only alternate strains labeled for readability. M, male; F, female. (**B**) Strains ranked by saline infusions. (**C**) Cocaine and saline infusions, ranked by cocaine. P is significance of $H^2$ difference between cocaine and saline. (**D**) Scatterplot shows significant correlation between cocaine and saline infusions averaged by strain ($R^2 = 0.2$, $t[1,82] = 4.5$, $P = 2.3 \times 10^{-5}$). Means ± s.e.m.*

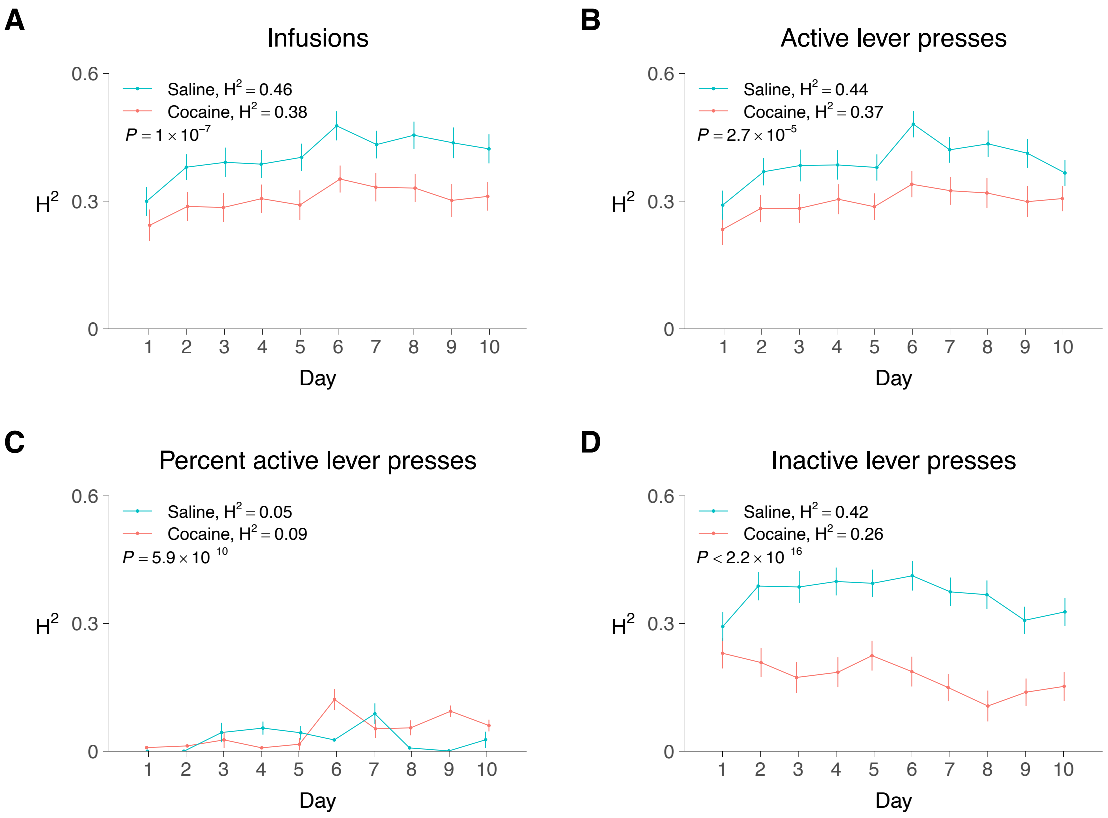


***Supplementary Figure S8.*** *Broad sense heritability ($H^2$) for cocaine and saline self-administration over 10 days. (**A**) Infusions (saline $H^2$ = 0.46 ± 0.001, $\chi^2$ = 2355, df = 1, P ≤ 2.2 × $10^{-16}$; cocaine $H^2$ = 0.38 ± 0.001, $\chi^2$ = 1756, df = 1, P ≤ 2.2 × $10^{-16}$; saline vs cocaine, P = 1.0 × $10^{-7}$). (**B**) Active lever presses (saline $H^2$ = 0.44 ± 0.002, $\chi^2$ = 2232, df = 1, P ≤ 2.2 × $10^{-16}$; cocaine $H^2$ = 0.37 ± 0.001, $\chi^2$ = 1739, df = 1, P ≤ 2.2 × $10^{-16}$; saline vs cocaine, P = 2.7 × $10^{-5}$). (**C**) Percent active lever presses (saline $H^2$ = 0.05 ± 0.001, $\chi^2$ = 109, df = 1, P ≤ 2.2 × $10^{-16}$; cocaine $H^2$ = 0.09 ± 0.002, $\chi^2$ = 257, df = 1, P ≤ 2.2 × $10^{-16}$; saline vs cocaine, P = 5.9 × $10^{-10}$). (**D**) Inactive lever presses (saline $H^2$ = 0.42 ± 0.001, $\chi^2$ = 2094, df = 1, P ≤ 2.2 × $10^{-16}$; cocaine $H^2$ = 0.26 ± 0.002, $\chi^2$ = 993, df = 1, P ≤ 2.2 × $10^{-16}$; saline vs cocaine, P ≤ 2.2 × $10^{-16}$). Saline vs. cocaine P values shown in graphs (sampling without replacement). Means ± s.e.m.*

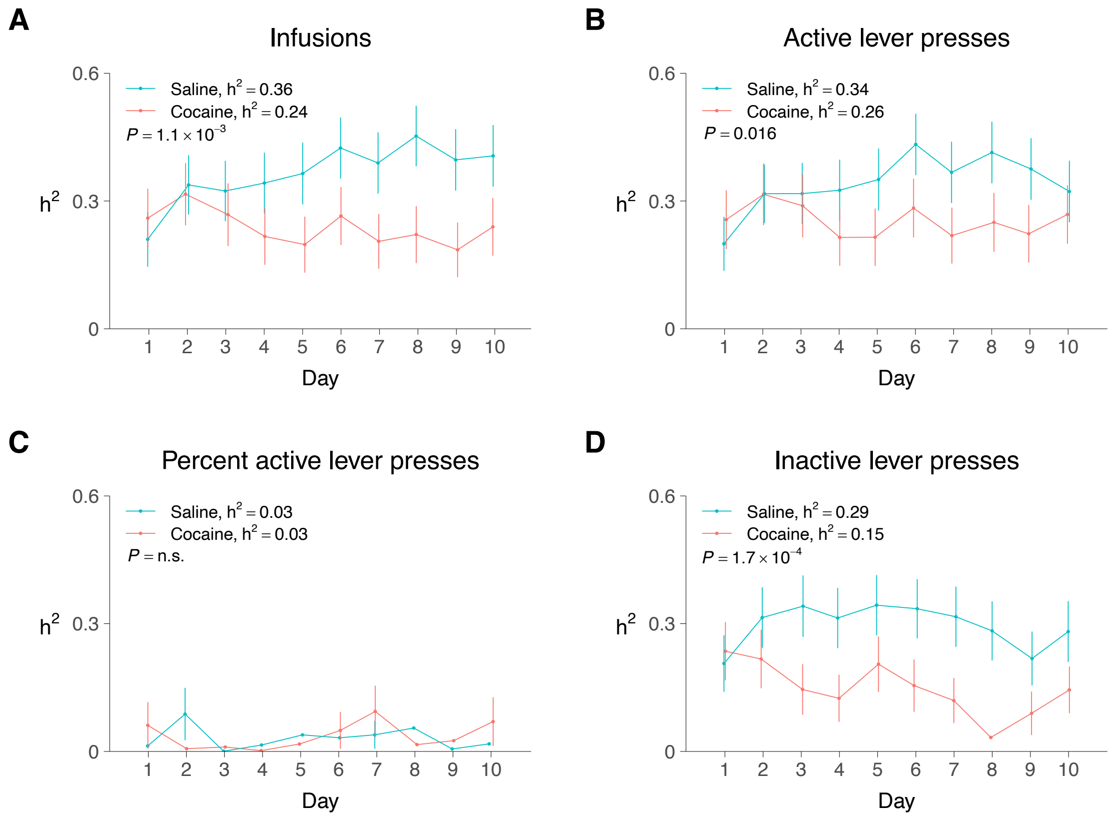


***Supplementary Figure S9.*** *Additive heritability ($h^2$) for cocaine and saline self-administration over 10 days. (**A**) Infusions (saline $h^2$ = 0.36 ± 0.02, t[1,9] = 17.0, P = 3.8 × $10^{-8}$, one sample t-test; cocaine $h^2$ = 0.24 ± 0.01, t[1,9] = 18.8, P = 1.6 × $10^{-8}$; saline vs cocaine, $\chi^2$ = 10.7, df = 1, P =1.1 × $10^{-3}$). (**B**) Active lever presses (saline $h^2$ = 0.34 ± 0.02, t[1,9] = 16.8, P = 4.3 × $10^{-8}$; cocaine $h^2$ = 0.26 ± 0.01, t[1,9] = 22.5, P = 3.2 × $10^{-9}$; saline vs cocaine, $\chi^2$ = 5.8, df = 1, P =0.016). (**C**) Percent active lever presses (saline $h^2$ = 0.03 ± 0.01, t[1,9] = 3.7, P = 5.2 × $10^{-3}$; cocaine $h^2$ = 0.03 ± 0.01, t[1,9] = 3.6, P = 6.3 × $10^{-3}$; saline vs cocaine, $\chi^2$ = 0, df = 1, P =1). (**D**) Inactive lever presses (saline $h^2$ = 0.29 ± 0.02, t[1,9] = 19.2, P = 1.3 × $10^{-8}$; cocaine $h^2$ = 0.15 ± 0.02, t[1,9] = 7.6, P = 3.4 × $10^{-5}$; saline vs cocaine, $\chi^2$ = 14.1, df = 1, P =1.7 × $10^{-4}$). Saline vs. cocaine P values shown in graphs. Means ± s.e.m.*

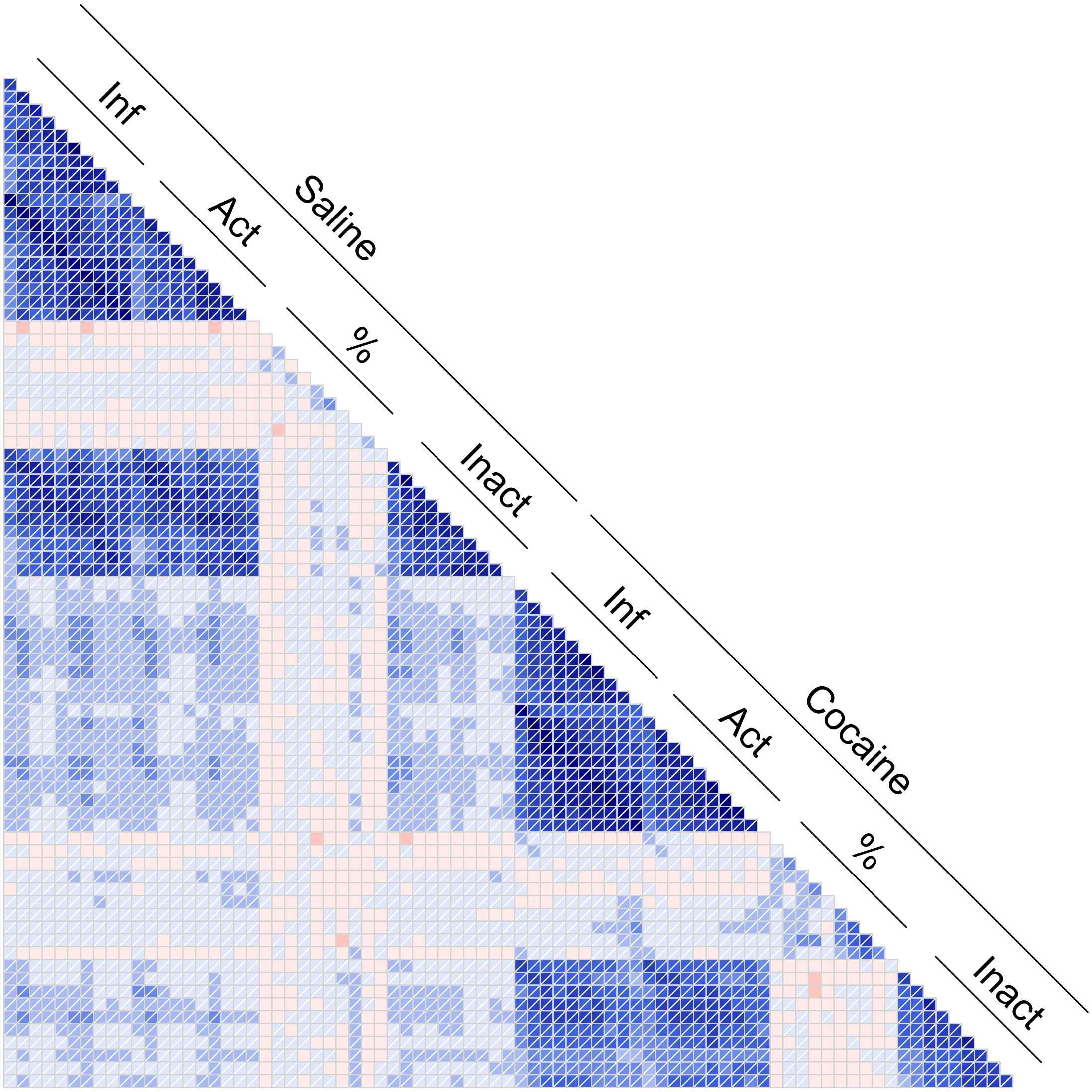


***Supplementary Figure S10.*** *Correlogram of -log$_{10}$P values from genome-wide association studies (GWASs) for each of the ten days of saline or cocaine IVSA; infusions, active lever presses, percent active lever presses and inactive lever presses.*

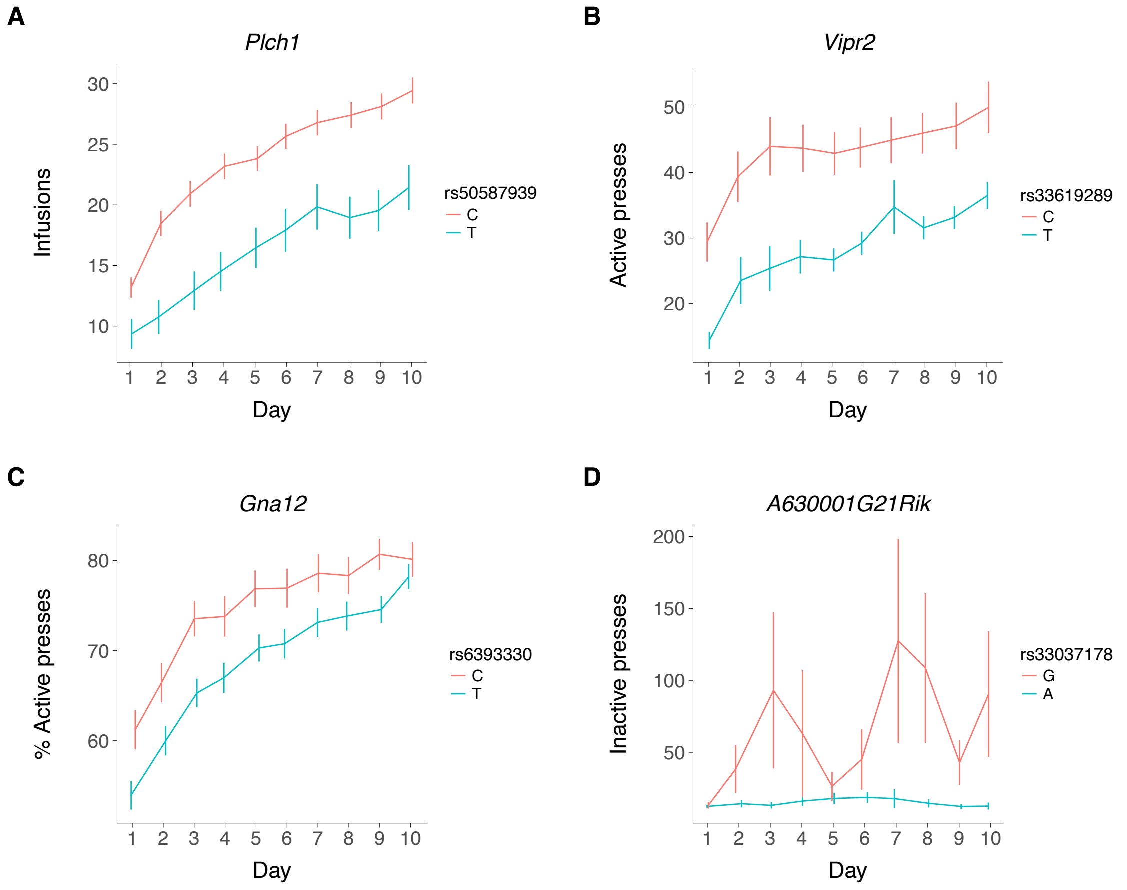


***Supplementary Figure S11.*** *Effects on cocaine IVSA of selected loci from longitudinal genome scans. (**A**) Infusions. (**B**) Active lever presses. (**C**) Percent active lever presses. (**D**) Inactive presses.*

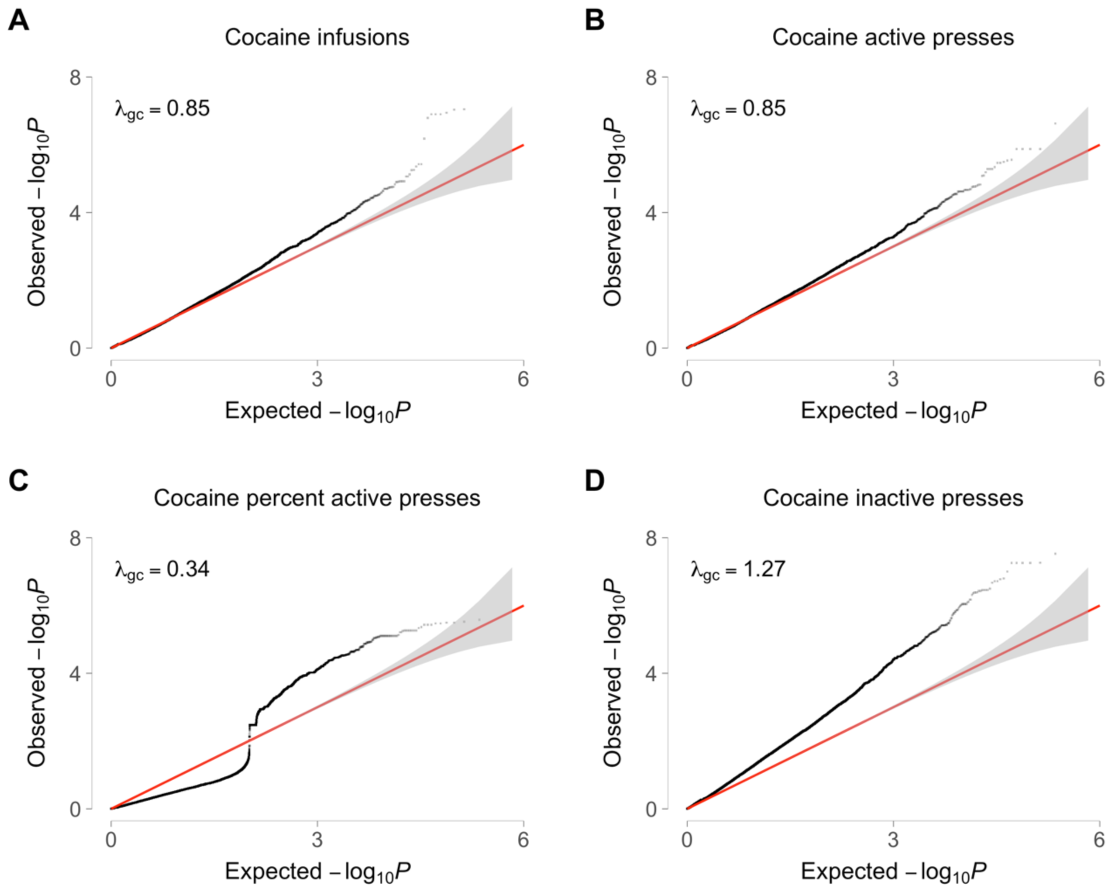


***Supplementary Figure S12.*** *Quantile-quantile (QQ) plots from longitudinal genome scans. (**A**) Infusions. (**B**) Active lever presses. (**C**) Percent active lever presses. (**D**) Inactive lever presses. $\lambda_{gc}$, genomic inflation factor.*

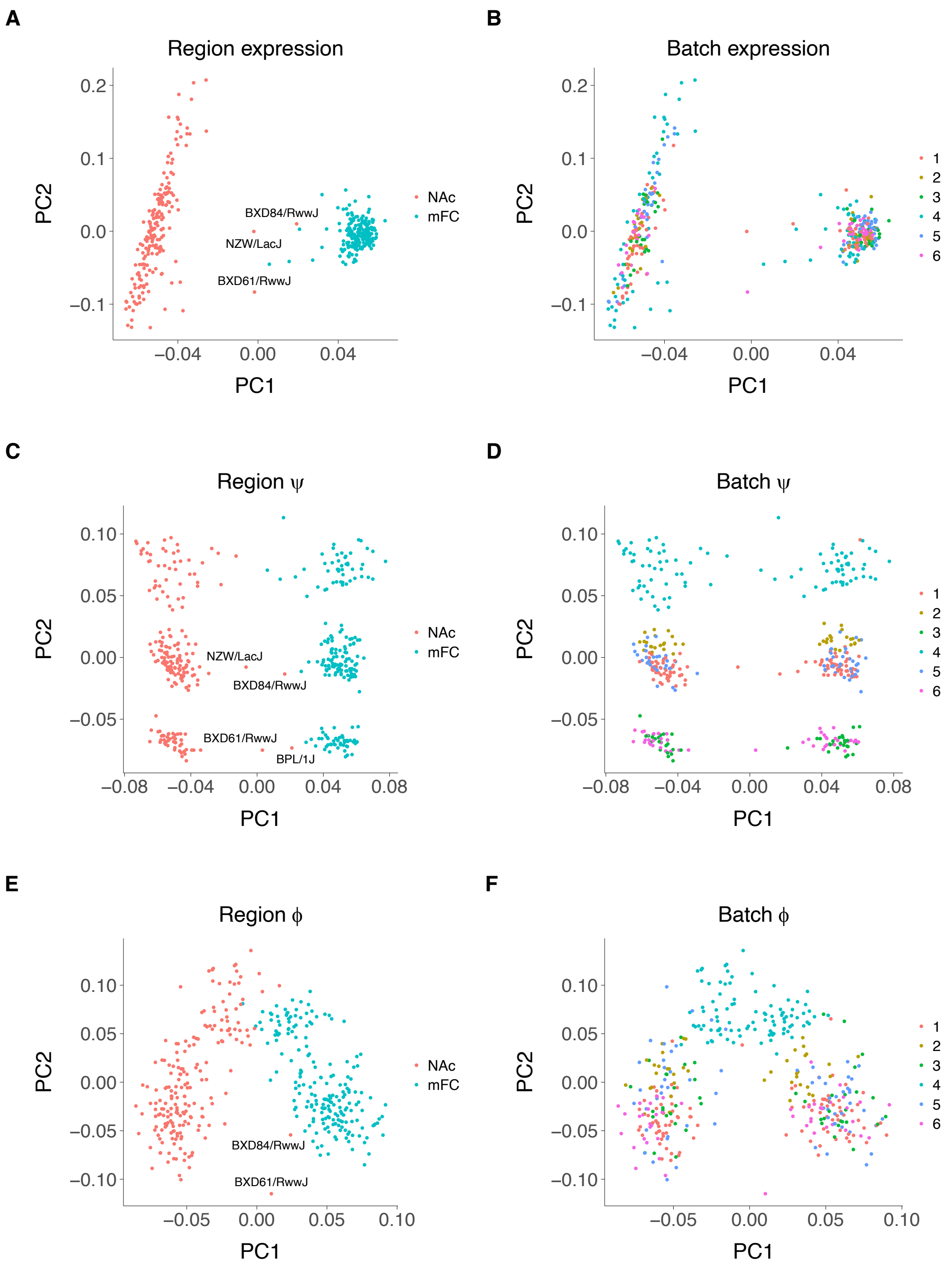


***Supplementary Figure S13.*** *Principal components analysis (PCA) of RNA-Seq. (**A**) Expression separated by region. (**B**) Expression separated by batch. (**C**) Splicing (ψ) separated by region. (**D**) Splicing separated by batch. (**E**) Editing (ϕ) separated by region. (**F**) Editing separated by batch. Potentially misassigned samples labeled with strain names. All potentially misassigned samples have a correctly assigned sample of the same strain but opposite sex.*

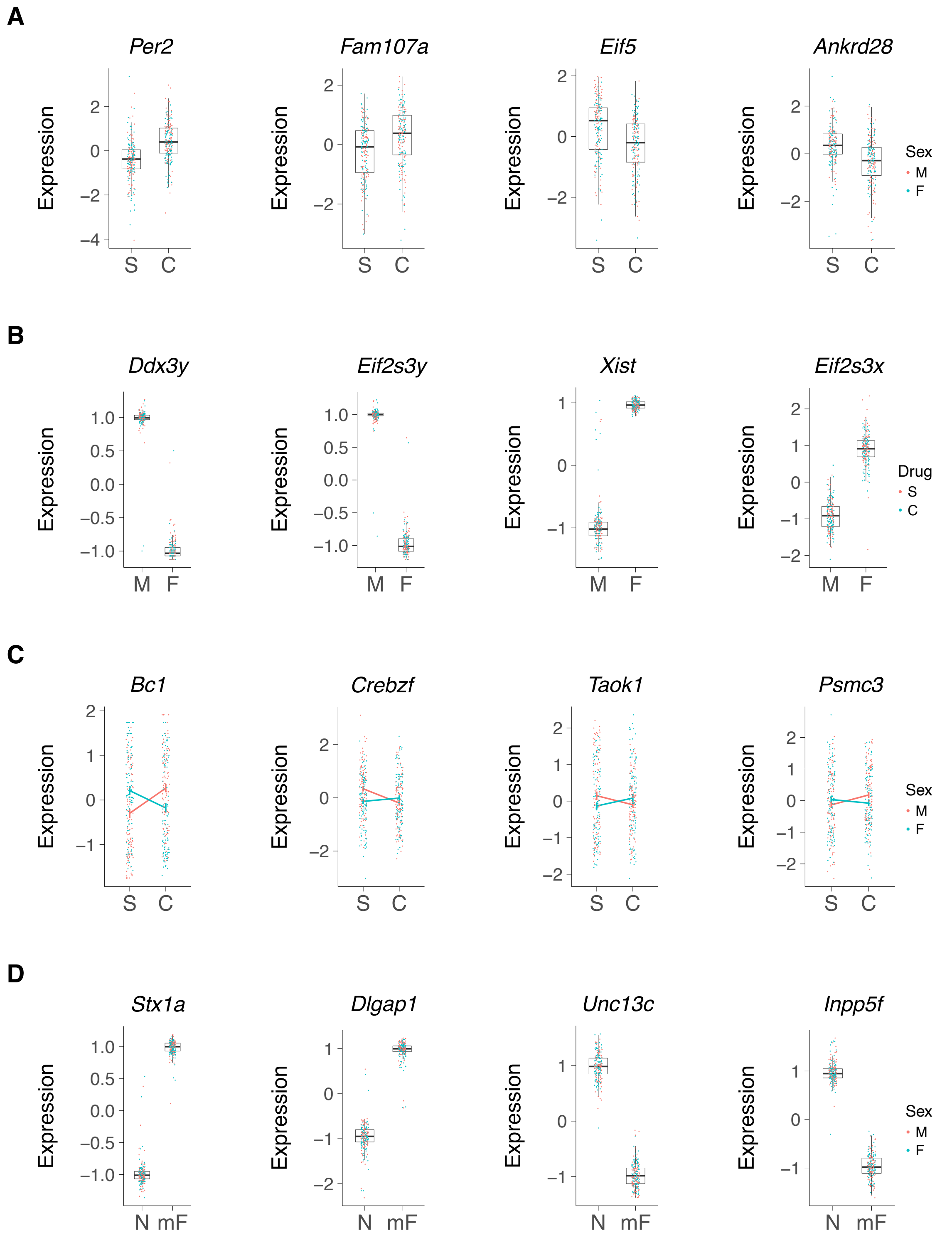


***Supplementary Figure S14.*** *Significantly regulated transcripts. (**A**) Transcripts regulated by infusate. S, saline; C, cocaine. (**B**) Sex. M, male; F, female. (**C**) Sex/infusate interaction. (**D**) Region. N, NAc (nucleus accumbens); mF, mFC (medial frontal cortex). FDR < 0.05.*

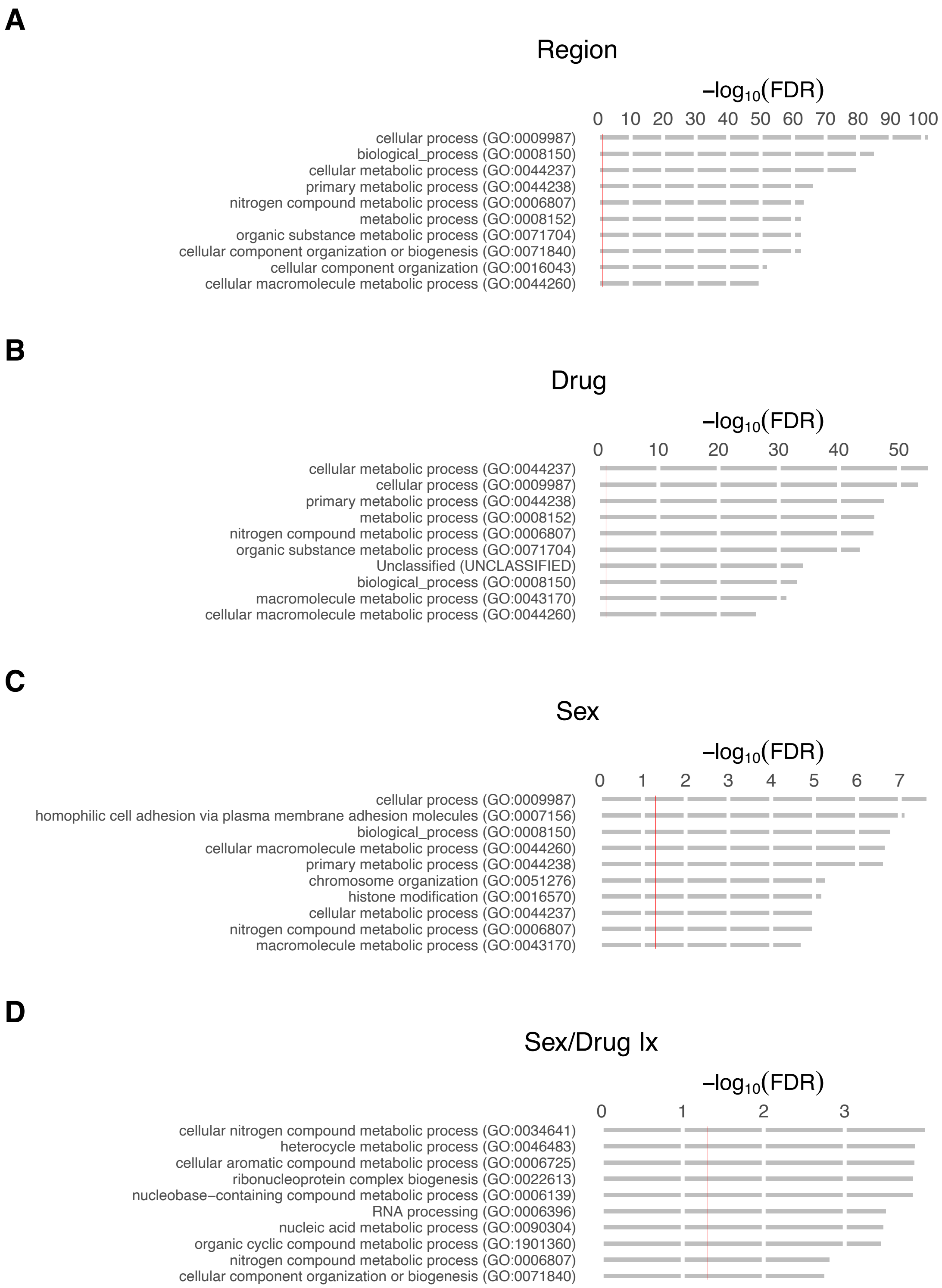


***Supplementary Figure S15.*** *Gene ontology (GO) analysis of transcript abundance. Biological process. Regulated genes with FDR < 0.05 chosen for analysis. Red lines, GO FDR = 0.05.*

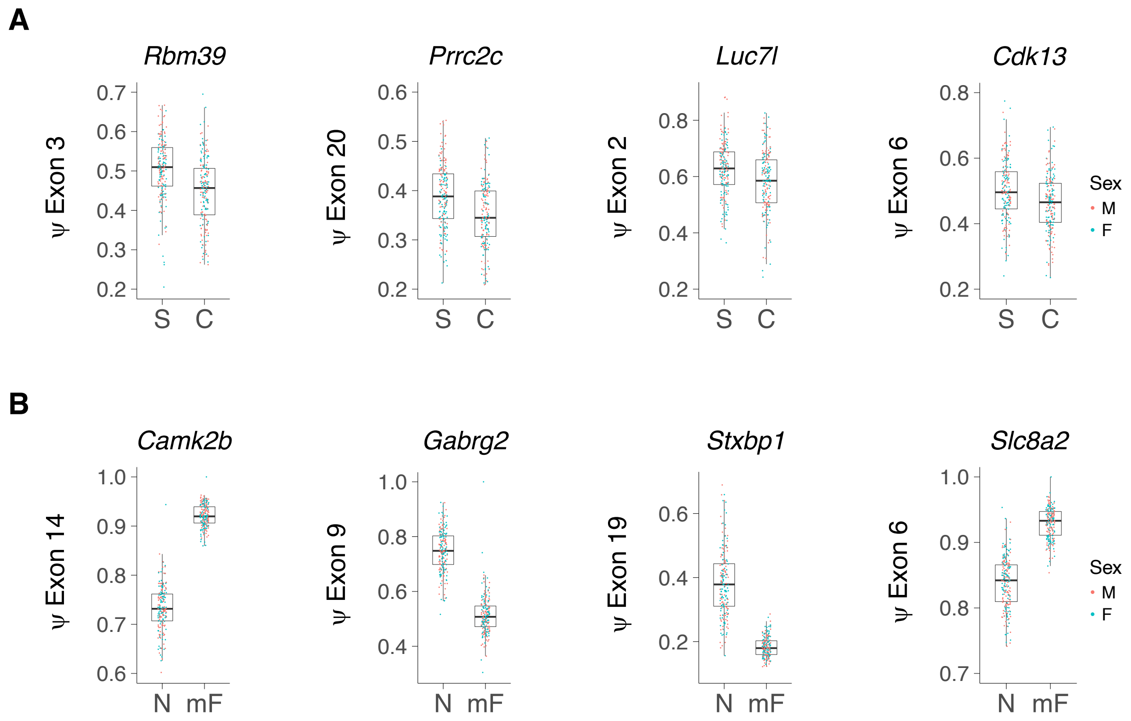


***Supplementary Figure S16.*** *Significantly regulated spliceforms. (**A**) Spliceforms regulated by infusate. ψ, percent spliced in or psi. S, saline; C, cocaine. (**B**) Region. N, NAc; mF, mFC. M, male; F, female. FDR < 0.05.*

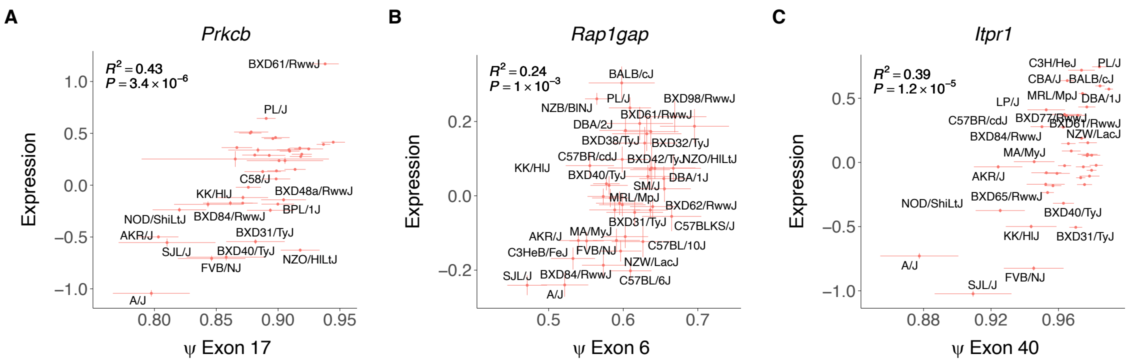


***Supplementary Figure S17.*** *Spliceforms affect RNA abundance. (**A**) Prkcb. (**B**) Rap1gap. (**C**) Itpr1. Percent spliced in (ψ). Strain means ±s.e.m. $R^2$ and P values are strain averaged results, linear mixed model FDRs < $2.2 \times 10^{-16}$.*

**A**

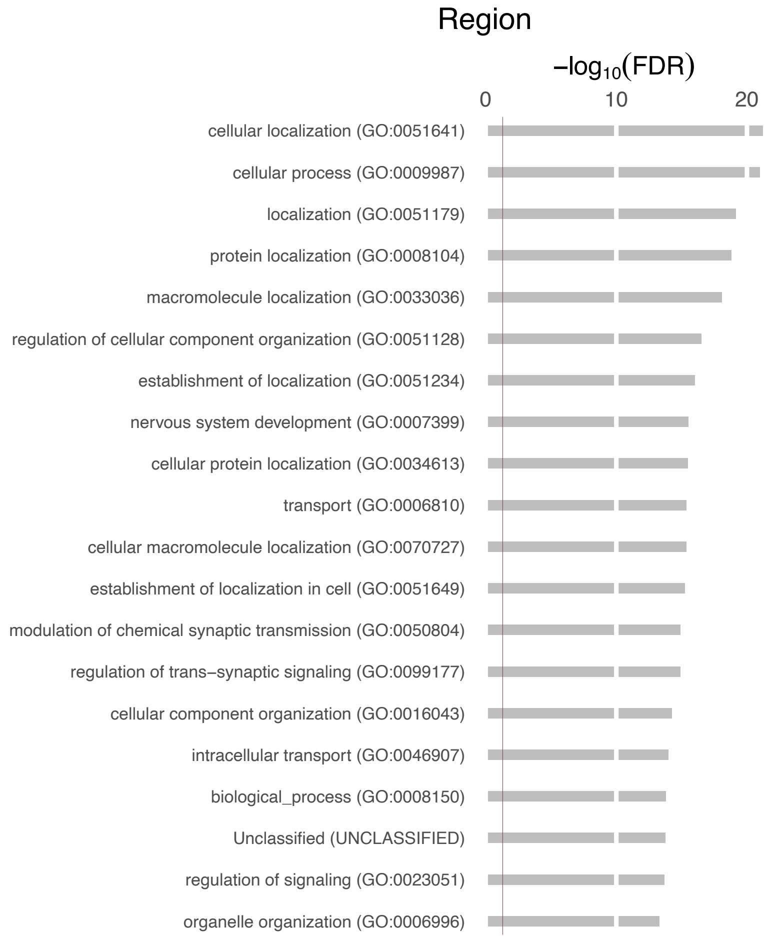


**B**

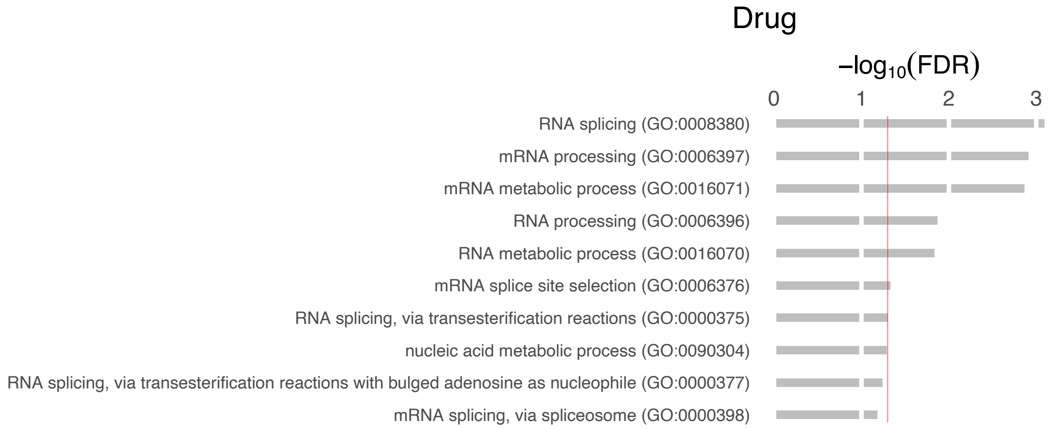


***Supplementary Figure S18.*** *Gene ontology (GO) analysis of splicing. Biological process. Regulated spliceforms with FDR < 0.05 chosen for analysis. Red lines, GO FDR = 0.05.*

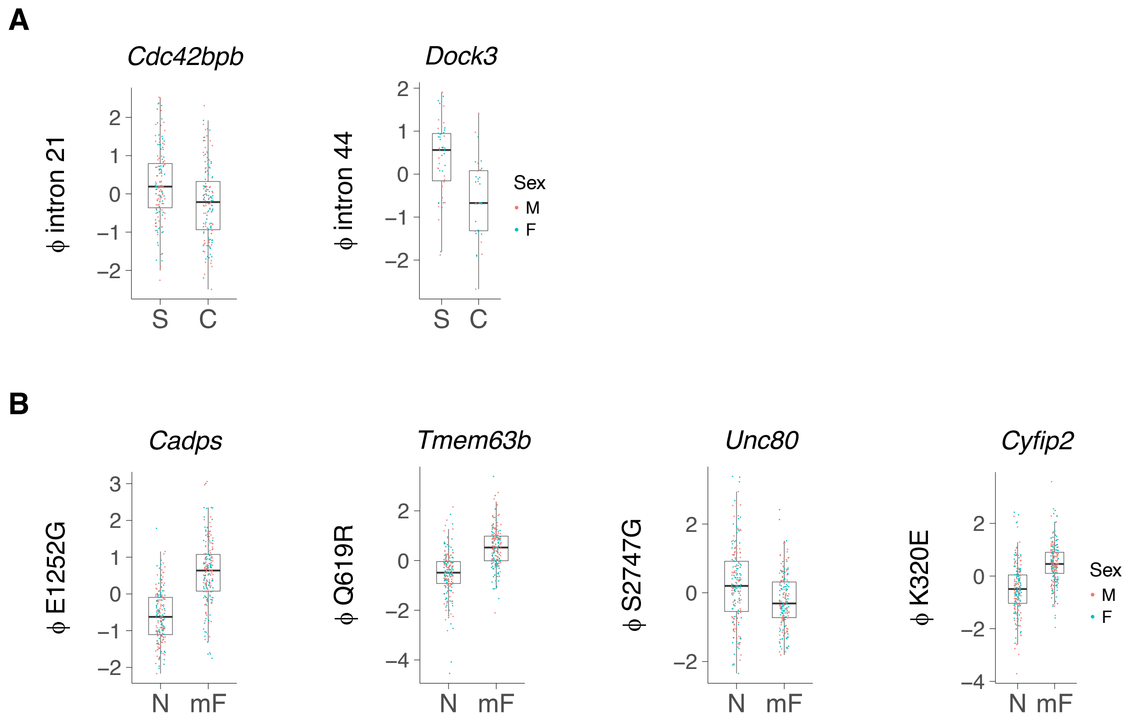


***Supplementary Figure S19.*** *Significantly regulated RNA editing. (**A**) RNA editing regulated by infusate. Both editing events are in intronic Alu elements, B1_Mus2 (Cdc42bpb: chr12: 111309987), B1_Mur1 (Dock3: chr9: 106905884). ϕ, RNA editing. S, saline; C, cocaine. M, male; F, female. (**B**) Non-synonymous RNA editing sites regulated by brain region. Cadps: NM_001042617: exon29: c.A3755G: p.E1252G; Tmem63b: NM_198167: exon20: c.A1856G: p.Q619R; Unc80: NM_001368824: exon54: c.A8239G: p.S2747G; Cyfip2: NM_133769: exon10 :c.A958G: p.K320E. N, NAc; mF, mFC. FDR < 0.05.*

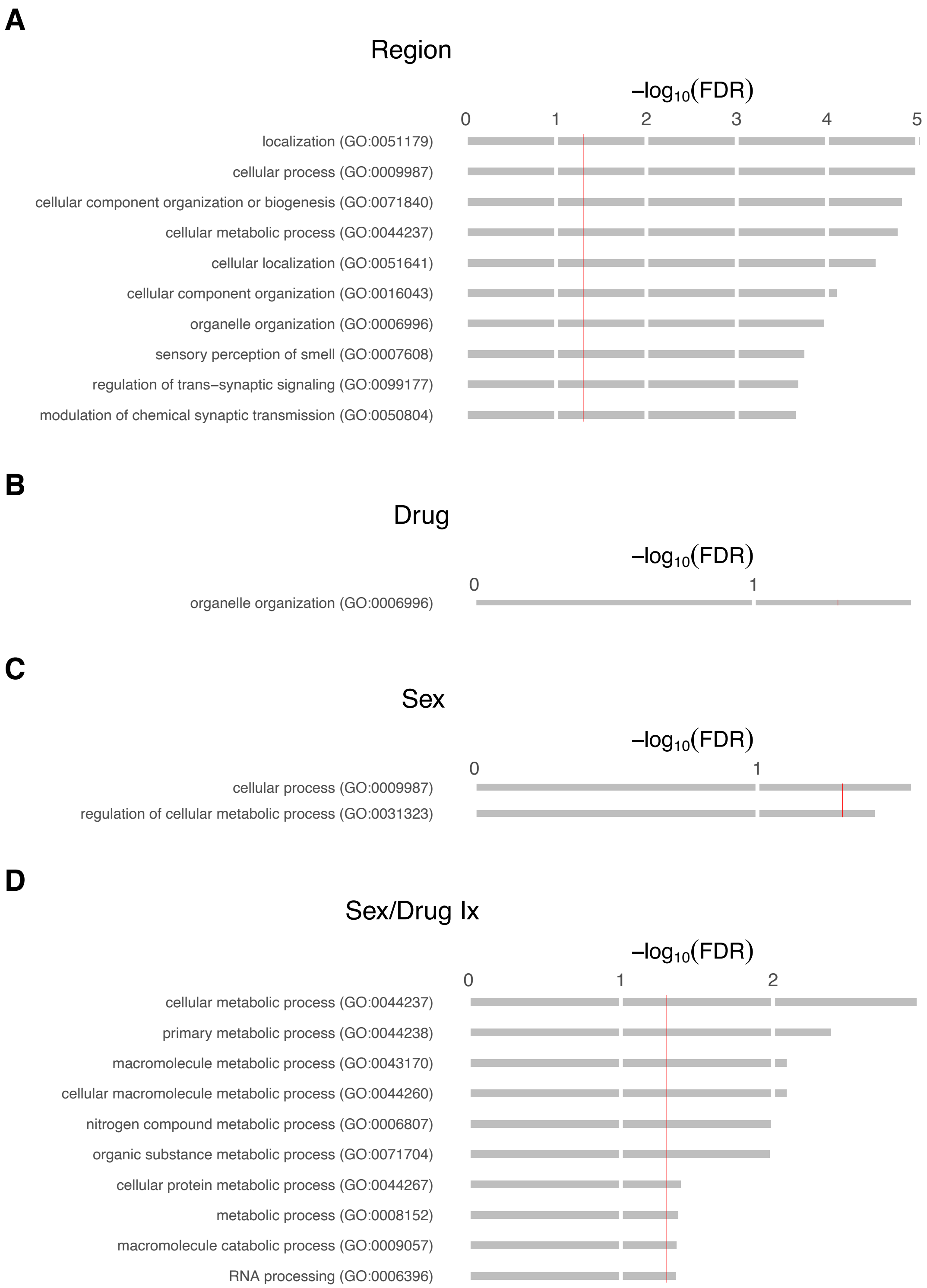


***Supplementary Figure S20.*** *Gene ontology (GO) analysis of RNA editing. Biological process. Regulated editing sites with P < 0.05 chosen for analysis. Red lines, GO FDR = 0.05. Organelle organization in (**B**), FDR < 0.05 (difficult to see red line).*

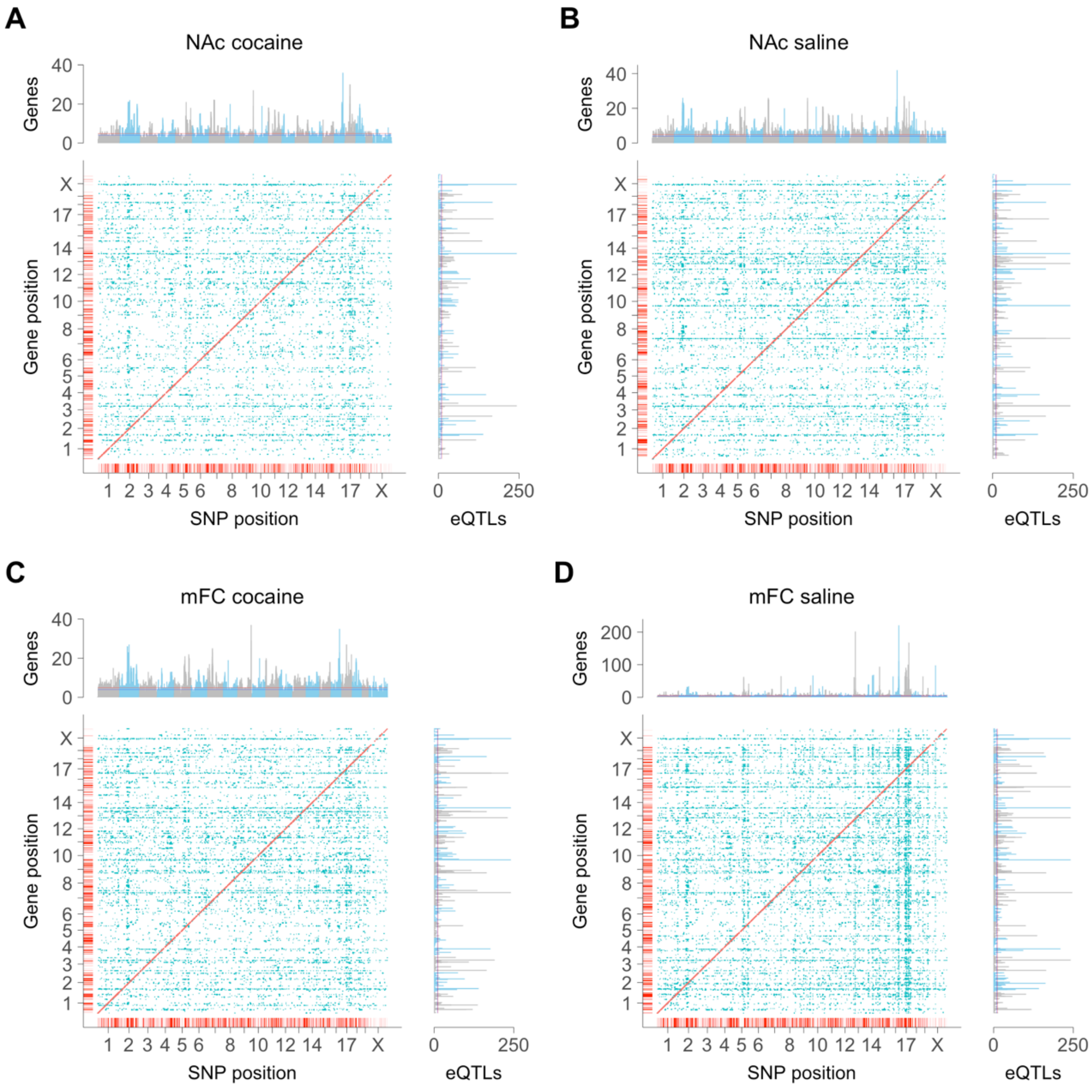


***Supplementary Figure S21.*** *Cis and trans expression quantitative trait loci (eQTLs) for transcript abundance. (**A**) NAc cocaine. Number of cis eQTLs (red), 4,469; trans eQTLs (blue), 10,463. (**B**) NAc saline. Number of cis eQTLs, 4,699; trans eQTLs, 13,354. (**C**) mFC cocaine. Number of cis eQTLs, 5,152; trans eQTLs, 14,609. (**D**) mFC saline. Number of cis eQTLs, 5,057; trans eQTLs, 18,091. Marginal plots, blue lines, FDR < 0.05; red lines, FDR < 0.01 (Poisson).*

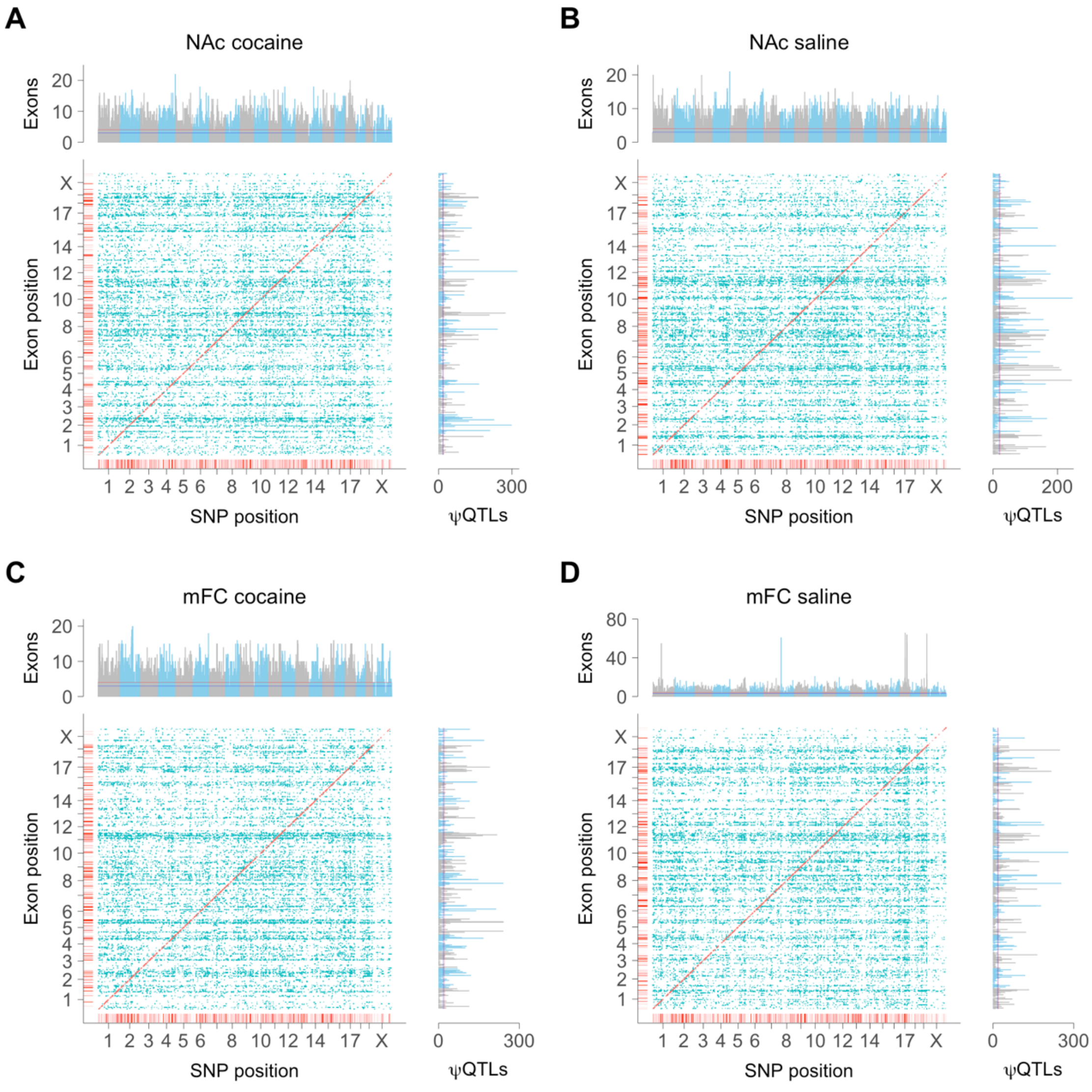


***Supplementary Figure S22.*** *Cis and trans splicing QTLs (ψQTLs). (**A**) NAc cocaine. Number of cis ψQTLs (red), 1,398; trans ψQTLs (blue), 25,140. (**B**) NAc saline. Number of cis ψQTLs, 1,385; trans ψQTLs, 26,276. (**C**) mFC cocaine. Number of cis ψQTLs, 1,469; trans ψQTLs, 25,532. (**D**) mFC saline. Number of cis ψQTLs, 1,450; trans ψQTLs, 25,028. Percent spliced in, ψ. Marginal plots, blue lines, FDR < 0.05; red lines, FDR < 0.01 (Poisson).*

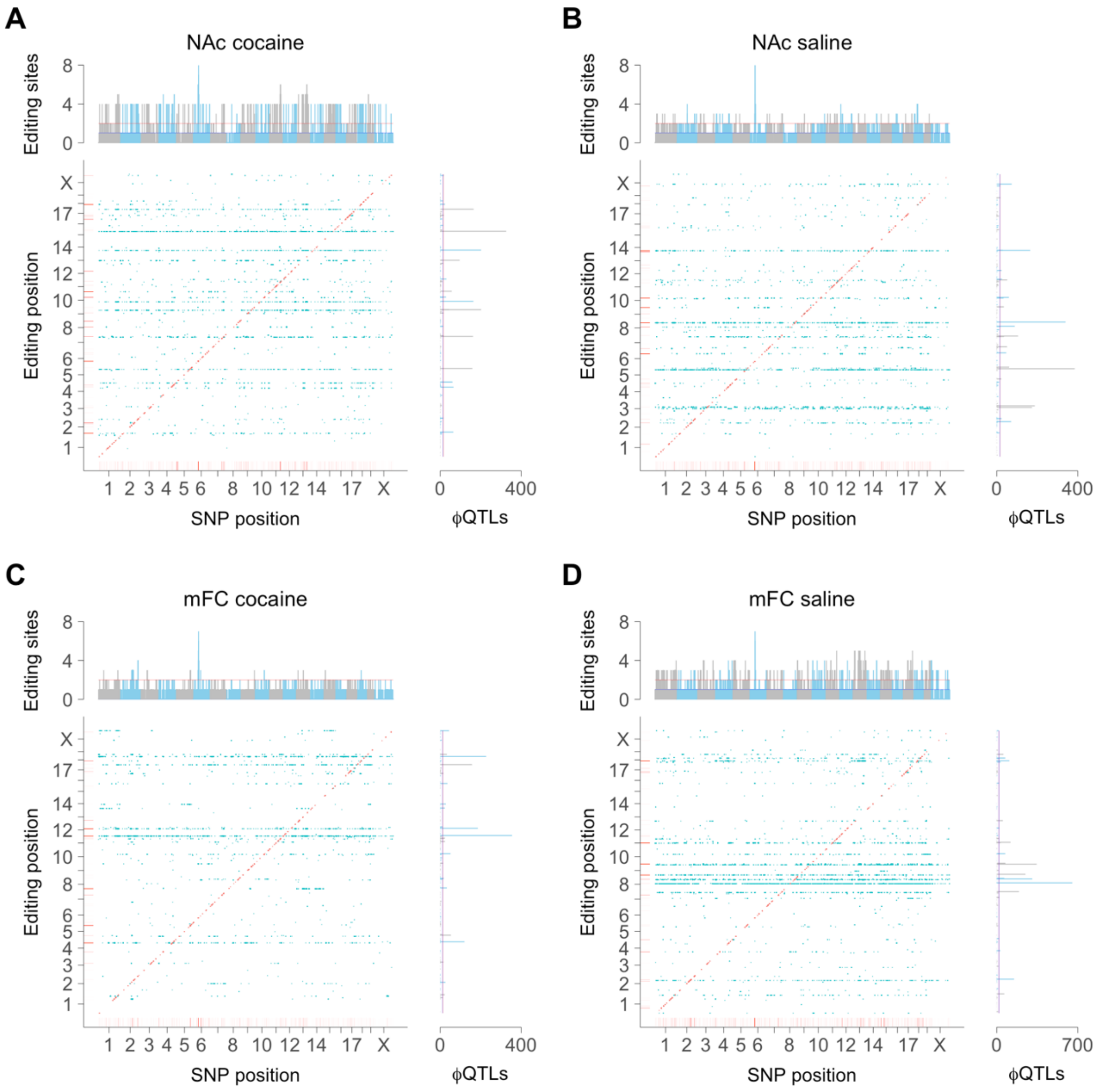


***Supplementary Figure S23.*** *Cis and trans editing QTLs (ϕQTLs). (**A**) NAc cocaine. Number of cis ϕQTLs (red), 290; trans ϕQTLs (blue), 2,507; editing ascertainment rate, 36 ± 0.3% of samples. (**B**) NAc saline. Number of cis ϕQTLs, 284; trans ϕQTLs, 2,614; ascertainment, 36 ± 0.3%. (**C**) mFC cocaine. Number of cis ϕQTLs, 274; trans ϕQTLs, 1,880; ascertainment, 37 ± 0.3%. (**D**) mFC saline. Number of cis ϕQTLs, 240; trans ϕQTLs, 3,301; ascertainment, 37 ± 0.3%. Editing ratio, ϕ. Marginal plots, blue lines, FDR < 0.05, red lines, FDR < 0.01 (Poisson).*

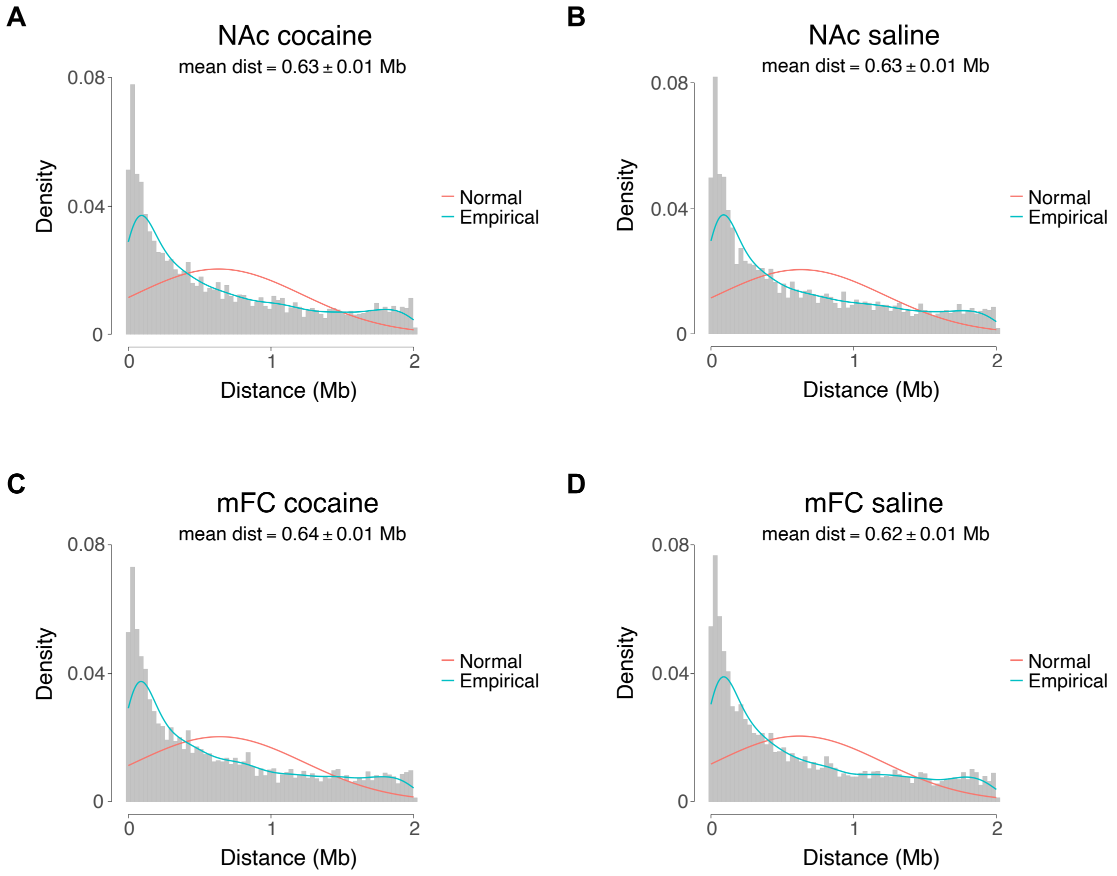


***Supplementary Figure S24.*** *Distances between cis eQTLs and the corresponding genes. (**A**) NAc cocaine. (**B**) NAc saline. (**C**) mFC cocaine. (**D**) mFC saline. Red line, normal distribution; turquoise line, empirical distribution.*

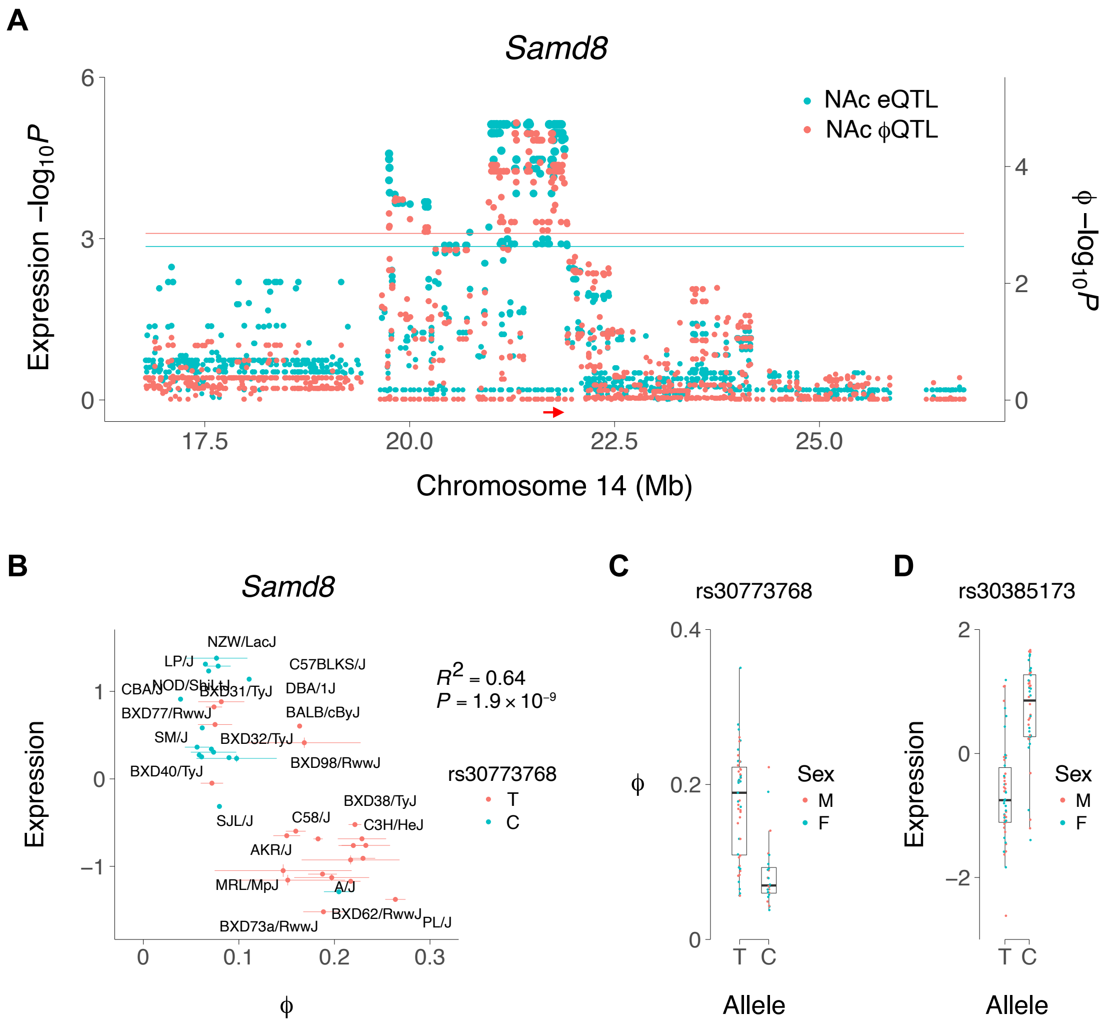


***Supplementary Figure S25.*** *RNA editing of Samd8 in NAc cocaine. (**A**) Coincident Samd8 cis editing QTL (ϕQTL) and cis expression QTL (eQTL). Editing location at 21,797,711 bp on Chromosome 14 in B1_Mm Alu element in the 3′ untranslated region of Samd8. Red arrow, location of Samd8. Red and blue horizontal lines, respective significance thresholds. (**B**) Allele T of rs30773768, the SNP most strongly correlated with Samd8 editing, associated with higher Samd8 editing and lower expression. Means ± s.e.m. for each strain. (**C**) Allele effect of rs30773768 on ϕ of Samd8. Individual samples shown. (**D**) Allele effect of rs30385173, the SNP most strongly correlated with Samd8 expression. SNPs rs30385173 and rs30773768 are in linkage disequilibrium ($D' = 1$, $R^2 = 0.94$, $P < 2.2 \times 10^{-16}$).*

**A**

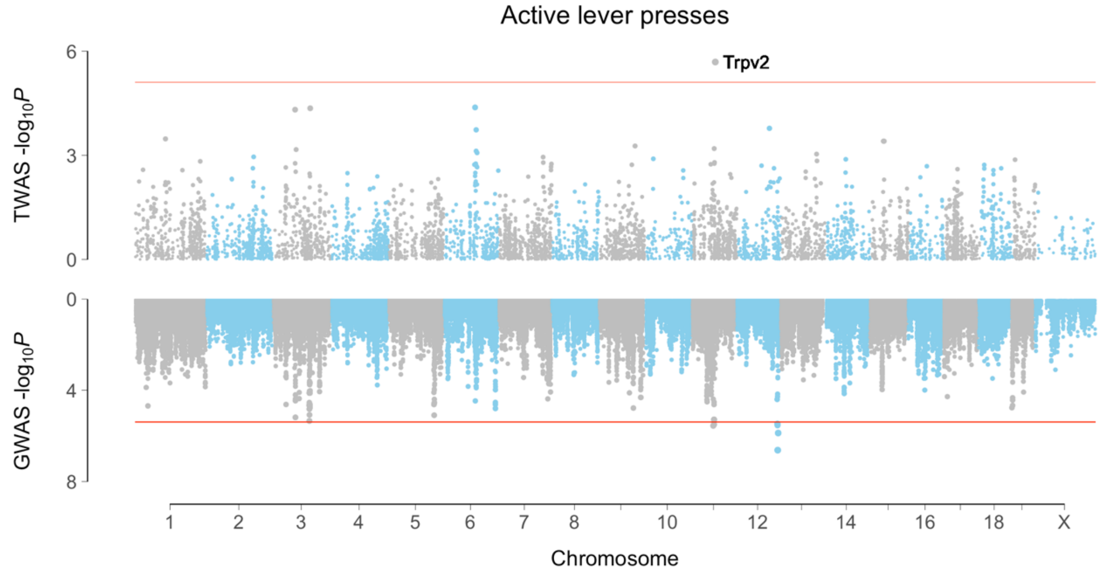


**B**

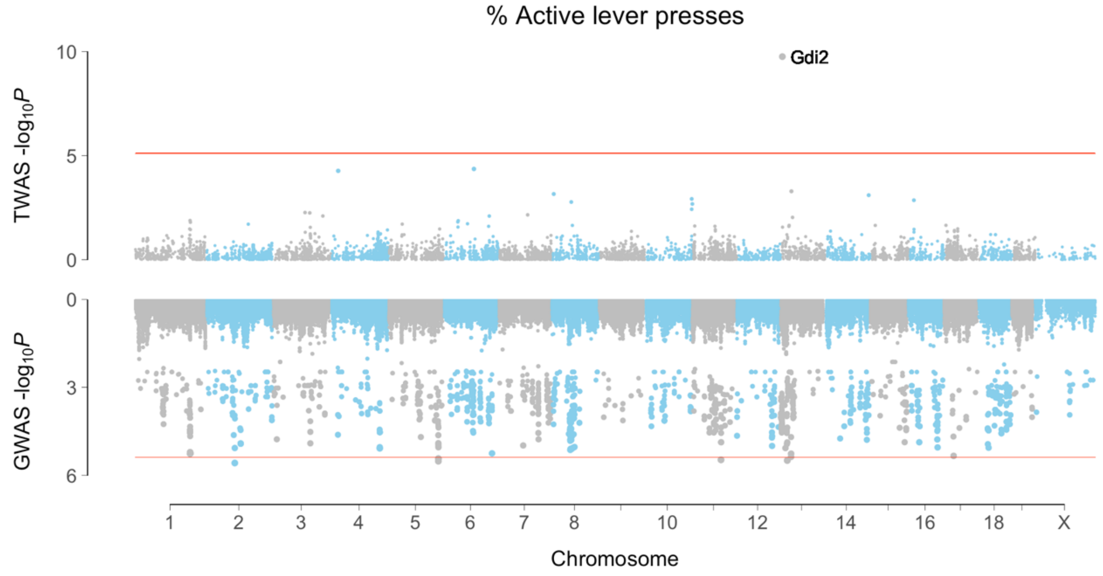


***Supplementary Figure S26.*** *Transcriptome-wide association studies (TWASs) of cocaine IVSA using RNA-Seq of NAc from drug-treated mice.* ***(A)*** *Active lever presses.* ***(B)*** *Percent active lever presses.*

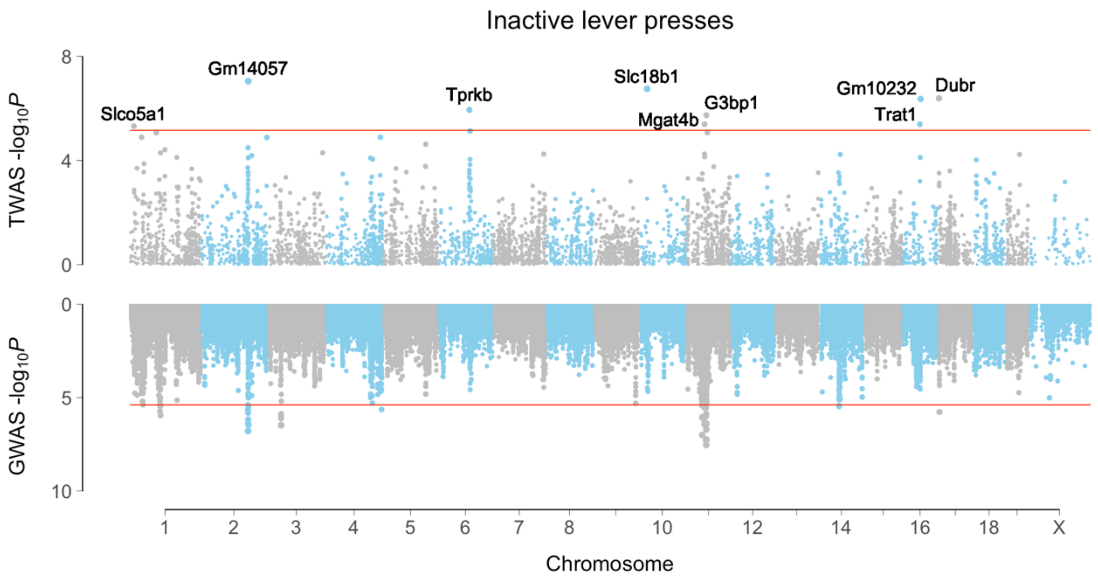


***Supplementary Figure S27.*** *TWAS of cocaine IVSA inactive lever presses using RNA-Seq of mFC from drug-treated mice.*
